# The differential mechanisms of eIF4A1-mediated translational activation instructed by distinct RNA features

**DOI:** 10.1101/2025.03.24.644895

**Authors:** Tobias Schmidt, Andrew P. Turnbull, Amit Gupta, Joseph A. Waldron, Katie Pollock, June Munro, Ridvan Nepravishta, Pauline Herviou, Adrianna Dąbrowska, Kelly Hodge, Allyson M. Voelker, Leon Pang, Camila Pasquali, Georgios Kanellos, Sara Zanivan, Owen J. Sansom, Jean-Denis Beaudoin, Martin Bushell

## Abstract

All mRNAs require eukaryotic translation initiation factor (eIF) 4A1 for translation through its different functions: loading of the pre-initiation complex onto mRNAs and unwinding of RNA structure. eIF4A1 is the catalytic subunit of the cap-binding eIF4F complex and presumed to select the mRNAs for translation through these activities, instructed by signalling pathways driving cell fate. The mechanisms underlying activation of the distinct eIF4A1 functions and establish translational selectivity are unknown.

Here, we have unravelled the complexity of mRNA selection by eIF4A1. We have mechanistically characterised the biological and atomic basis of inhibition by gain- and loss-of-function eIF4A1-inhibitors eFT226 and hippuristanol, and used machine learning to model the mRNA features associated with specific inhibition. This uncovered the eIF4A1 function – mRNA sequence relationship: 5’UTRs containing C/CG-rich require efficient mRNA loading by eIF4F which is specifically targeted by hippuristanol, while 5’UTRs harbouring alternatives starts sites together with AG-rich motifs utilise eIF4A1 for start site selection, specifically perturbed by eFT226. Our model is validated through a massively parallel reporter assay using 5’UTRs from a distinct evolutionary origin. This prompted us to examine the conservation between mRNA sequence and eIF4A1 function, and revealed their co-development.

Our findings highlight opportunities for novel therapeutic strategies targeting eIF4A1 and for improved design of mRNA-based therapeutics.

## Hippuristanol and eFT226 show distinct modes of action in cells

We have shown previously that eIF4A1 activity is regulated by specific RNA sequences *in vitro*^1^. Thus, we hypothesised that distinct mRNA sequence features activate different eIF4A1 functions on eIF4A1-dependent mRNAs. To examine this, we followed a two-fold experimental strategy: (1) To enable deconvolution of eIF4A1-dependent translation, we treated cells with distinct classes of eIF4A1 inhibitors, hippuristanol and eFT226 (EFT226, Zotatifin, **Fig. 1a**), and, subsequently, monitored changes in cellular mRNA translation by ribosome-profiling and real-time proteomics. (2) To increase stringency of the analysis, the experiments were performed in two biological systems, human breast cancer MCF7 cells and organoids derived from a colorectal cancer mouse model (*VillinCre*^ER^ *Apc^-^*^/-^, *Kras*^G12D/+^).

**Figure 1.**
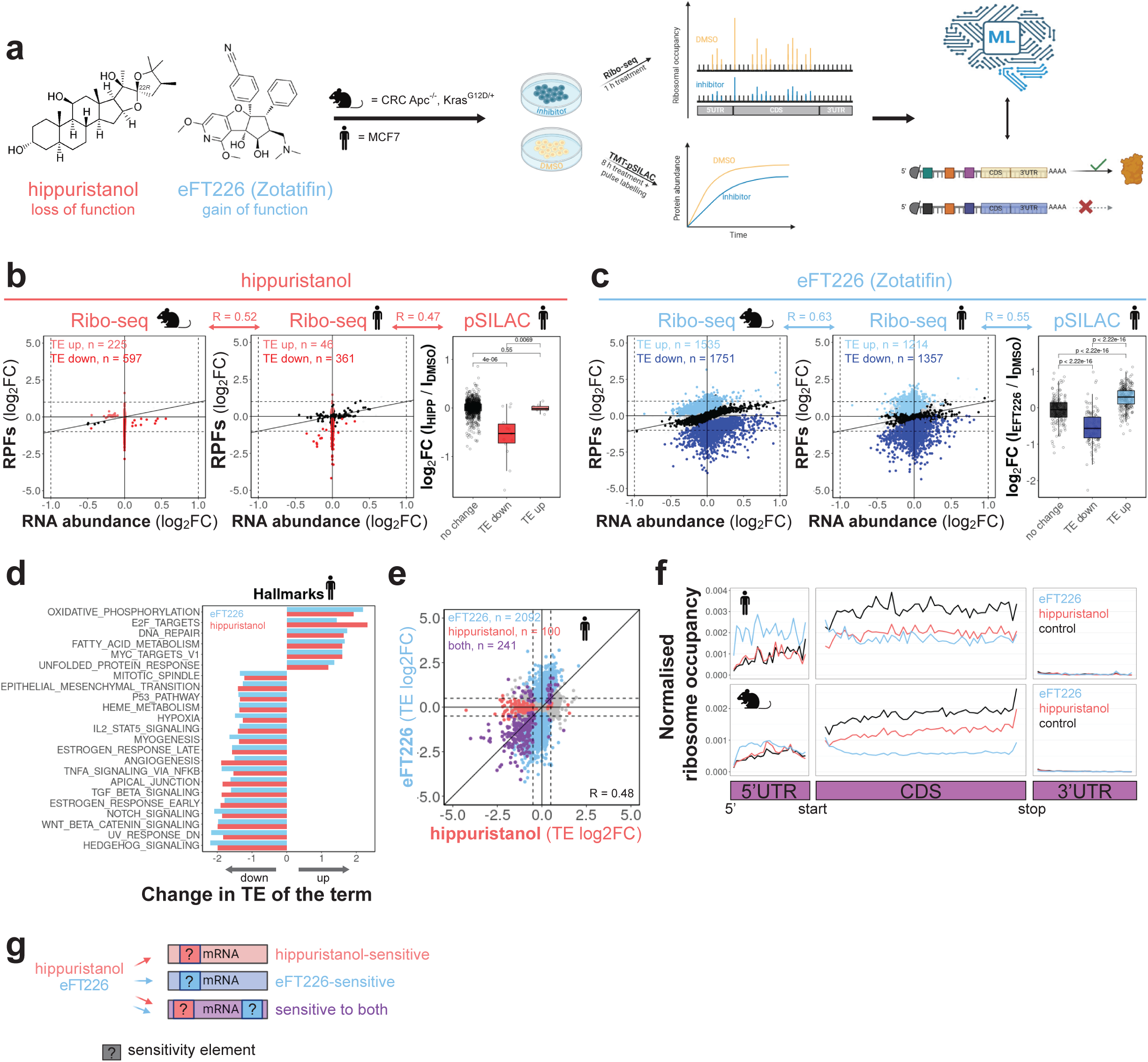
Hippuristanol and eFT226 show distinct modes of action in cells. a, Chemical structures of hippuristanol and eFT226 (Zotatifin), and experimental strategy to translationally profile eIF4A1-dependent mRNAs. Asterisk **b**, Ribo-seq (log2-fold change ribosomal occupancy), RNA-seq (log2-fold change RNA abundance) and TMT-pulsed SILAC (log2-fold change protein abundance) of genes after IC_50_ treatment of MCF7 and mouse organoids derived from a colorectal cancer model (Apc^-/-^, Kras^G12D/+^) with hippuristanol or **c**, eFT226. MCF7: Data are mean-per-gene from n repeat experiments. n(Ribo-seq) = 5, n(RNA-seq) = 6, n(pSILAC) = 4. mouse: Data are mean-per-gene from n biological repeats. n =3. P-values from Wilcoxon test. R – Pearson correlation coefficient. Box plot shows median ± interquartile range. **d**, Hallmark enrichment analysis based on changes in TE of genes following hippuristanol or eFT226 treatment of MCF7 cells. **e,** Mean log2-fold changes in translational efficiencies (TE) per gene following eFT226 and hippuristanol treatment in MCF7 cells. R – Pearson correlation coefficient **f**, Metagene analysis of the mean ribosome occupancy across the whole mRNA of all genes sensitive to both (purple group) compounds. **g**, Schematic presentation of the hypothesis that mRNAs contain specific sensitivity elements rendering them distinctly responsive to compound treatment.

Elaborating on points above, eIF4A1 is the pharmacological target of hippuristanol and eFT226^2,3^, however, their cellular modes of action are not fully characterised. *In vitro*, they induce distinct modes of action^2,4^, by which eFT226 follows a gain of function mechanism, while hippuristanol induces a loss of eIF4A1 function. Specifically, eFT226 improves clamping of eIF4A1 RNA-binding (**Supplementary Fig. 1a-c**)^4^, and, through this, stimulates multimerisation of eIF4A1 specifically on AG-rich sequences which enhances eIF4A1 unwinding (**Supplementary Fig. 1d-f**). In contrast, hippuristanol triggers RNA release leading to destabilised RNA-eIF4A1 complexes (**Supplementary Fig. 1a-c**) and thus inhibited RNA unwinding (**Supplementary Fig. 1d-f**) ^1,2^. Additionally, previous observations suggest that both compounds target eIF4A1 that is or is not incorporated into eIF4F distinctly^1,5^. Altogether, we, thus, hypothesised that the compounds perturb dissimilar eIF4A1 functions in cells. Hence, we wanted to directly compare and determine the cellular impact of these inhibitors with such different modes of actions (**Fig. 1a**).

We performed compound treatment in human and mouse systems as their eIF4A1 proteins are identical and basic mRNA sequence compositions of their transcriptomes correlate well (e.g., CDS composition) but also show dissimilarities (e.g., 5’UTR composition, **Supplementary Fig. 1g**). Thus, we expected that data integration from both species, would uncover more common and conserved characteristics of mRNA sequences involved in eIF4A1 functions.

MCF7 cells and mouse organoids were treated for one hour with IC_50_ concentrations of the compounds (**Supplementary Fig. 2a-b**) and changes in ribosome protected footprints (RPF) together with mRNA abundance quantified following standard sequencing and data processing procedures (**Supplementary Fig. 2c-h**). Consistent between mouse and human cells, hippuristanol primarily *down*-regulated the translational efficiency (TE) of its mRNA targets by modulating RPF levels (**Fig. 1b**), while eFT226 induced significant changes to both the mRNA and RPF levels of a broad number of targets leading to *up* and *down*- regulated TEs (**Fig. 1c**). Additionally, 8h-pulsed SILAC under the same treatment conditions in MCF7 cells confirmed that compound-induced translational regulation of target mRNAs, both *up*- and *down*-regulation, directly impacts protein output of these mRNAs in the same direction (**Fig. 1b-c, Supplementary Fig. 2i**). Hallmark and pathway enrichment analyses, exemplified with the human data, identified common functional mRNA families targeted by both compounds, including known eIF4A1-dependent pathways such as WNT signalling and regulation of cell cycle components (**Fig. 1d, Supplementary Fig. 2j**)^1,6–8^. Together, this approach allowed us to identify different classes of eIF4A1-dependent translation.

Interestingly, correlation between the changes in translational efficiencies following hippuristanol- and eFT226-treatment was limited (R_human_ = 0.48, R_mouse_ = 0.42, **Fig. 1e, Supplementary Fig. 2k**), revealing also pools of mRNAs that were only sensitive to either hippuristanol or eFT226 (**Fig. 1e, Supplementary Fig. 2l**). This was also true within mRNA families (**Supplementary Fig. 2m**). Strikingly, mRNAs targeted by either compound showed distinct changes in their 80S ribosomal occupancy within the 5’UTR and CDS of the mRNA (**Fig. 1f**). Specifically, eFT226-treatment resulted in an accumulation of ribosomes in the 5’UTR and a steep, gradual reduction of ribosomes along the CDS, while hippuristanol- treatment led to reduced ribosomal occupancy, equally across the CDS without changes to the 5’UTR (**Fig. 1f**). This clearly demonstrates that the two compounds perturbate translation through different modes of action affecting compound-specific mRNAs. This was consistent between mouse and human cells suggesting that different mRNA sequence elements are linked to specific eIF4A1 functions (**Fig. 1g)**.

## Hippuristanol and eFT226 target mRNAs with distinct 5’UTR and CDS features

To identify the most common mRNA features that are critical to predict compound sensitivity, and thus involved in eIF4A1 functions, we merged human and mouse data and applied gradient boosting, an unbiased machine learning process. By this, we aimed to computationally model the impact of mRNA features on the change in translational efficiency following individual compound treatment. For both eFT226 and hippuristanol, a selection of mRNA features, that correlated to varying degrees with the experimental data, was used for the computation (**Fig. 2a**). The resulting models were highly accurate and generated strong correlation with the experimental data (**Supplementary Fig. 3a**), while cross-predictions were substantially less accurate (**Supplementary Fig. 3b)**. The models identified 5’UTR length as the common and most influential mRNA feature predicting both hippuristanol and eFT226-sensitivity (**Supplementary Fig. 3c**), which is consistent with eIF4A1’s known role in mRNA translation^2,7,8^. To increase focus on other mRNA features, 5’UTR length was subsequently omitted from the models^9^, revealing frequency of 5’UTR purine-rich (AG- motifs) motifs and 5’UTR C/CG-content amongst the mRNA features predicting differential sensitivity to eFT226 or hippuristanol, respectively (**Fig. 2b-c, Supplementary Fig. 3d)**.

**Figure 2.**
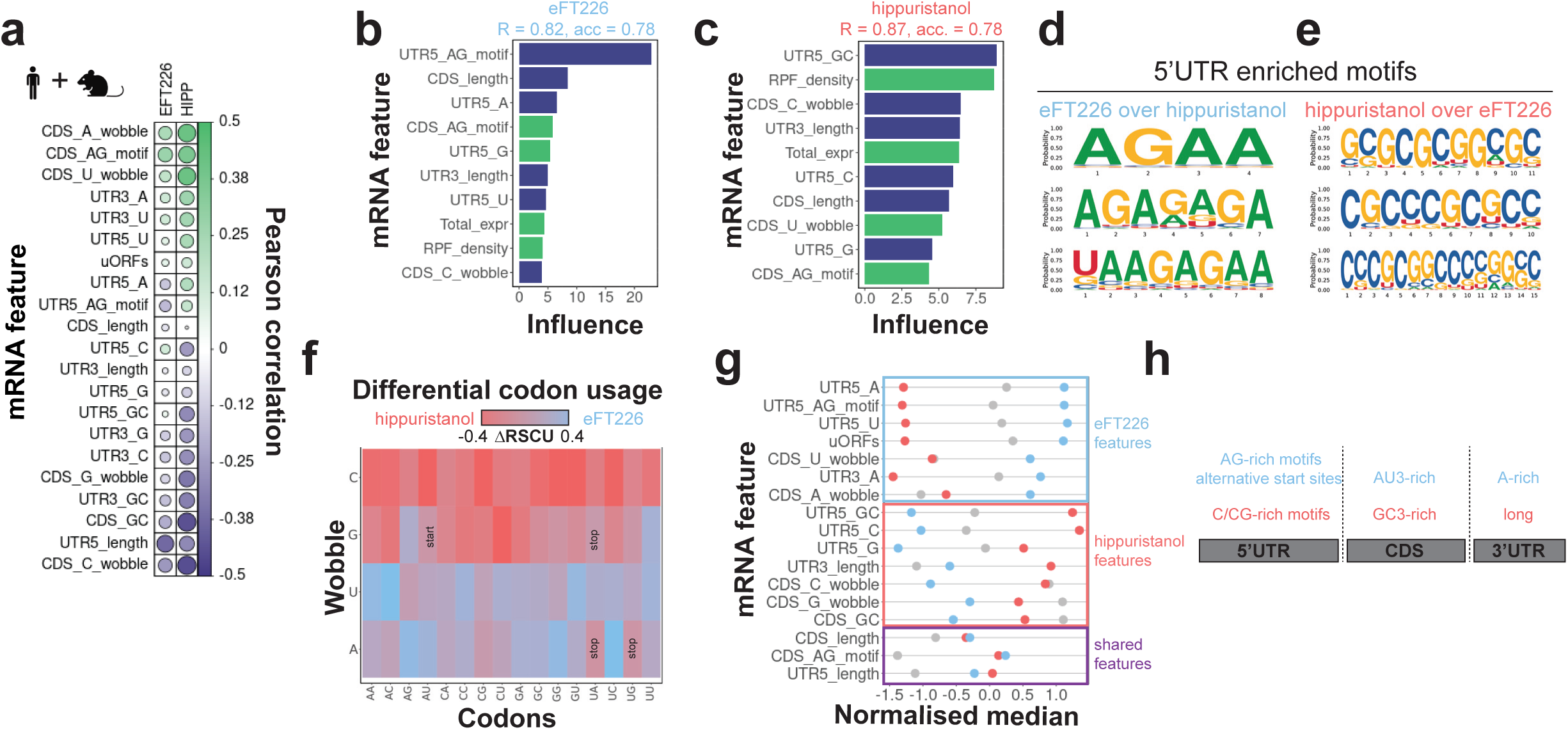
mRNA features drive differential sensitivity of mRNAs to eFT226 or hippuristanol treatment. a, Pearson correlation between mRNA features (in the format “region_type”) used in the gradient boosting and log2-fold changes in TE of mRNAs following hippuristanol or eFT226 treatment. Combined data from mouse and human cells **b**, Top ten mRNA features most influential in predicting mRNA sensitivity to eFT226 or **c**, Hippuristanol as derived by gradient boosting. Input data combined from human and mouse Ribo-seq. R – Pearson correlation between predicted and experimental log2-fold changes in TE (**Supplementary** Fig. 3d). Accuracy (acc) – Precision in predicting compound sensitivity tested again the input data. **d**, Top three differentially enriched 5’UTR sequence motifs of eFT226- or **e**, hippuristanol-sensitive mRNAs over the other one, respectively, for combined human and mouse mRNA targets. **f**, Differential synonymous codon usage of hippuristanol and eFT226-sensitive mRNAs. Data are mean-per-codon across all transcripts in the group. **g**, Per-feature normalised medians of mRNA features with significant difference (p-value < 0.1) between eFT226- (blue), hippuristanol-sensitive (red) and compound-insensitive mRNAs (grey). **h,** Cartoon presentation of mRNAs emphasising features most predictive for eFT226- or hippuristanol-sensitivity.

These features have been observed before in previous studies characterising rocaglamide and hippuristanol sensitive mRNAs separately^2,7,10^, validating our unbiased computational approach. Supporting this, we discovered distinct RNA sequence motifs differentially enriched in the 5’UTRs of eFT226 over hippuristanol mRNA targets and *vice versa* summarising their unique 5’UTR features (**Fig. 2d-e, Supplementary Fig. 3e**). These motifs represent distinctive modes of action and functions of eIF4A1^1^, with AG-motifs (eFT226 enriched) being related to increased clamping and multimerisation of eIF4A1 on AG-rich RNA elements^1,10^ (see also **Supplementary Fig. 1b-c**) , while C/CG-rich motifs (hippuristanol enriched) presumably representing stable RNA structures requiring unwinding activity of eIF4A1^1^.

To our surprise, the model also identified features of protein coding regions (CDS) of the target mRNAs including codon usage information as well as ribosome density and mRNA abundance, that influence sensitivity to hippuristanol and eFT226 (**Fig. 2b-c**).

Examination of the coding regions of target mRNAs revealed divergent preferences of codon usage (**Fig. 2f**) with eFT226 targets utilising A/U wobbles (AU3) while hippuristanol targets are enriched for G/C wobbles (GC3). In MCF7 cells this matched the codon bias of the highest (AU3-rich, top 10% of mRNAs) and lowest abundant mRNA (GC3-rich, bottom 10% of mRNAs), respectively, (**Supplementary Fig. 3f**), while this split was less clear in the highly proliferative mouse organoids (**Supplementary Fig. 3g**). Our data indicate that this may be related to a higher abundance of hippuristanol- than eFT226-sensitive mRNAs in this cancer model (**Supplementary Fig. 3h)**.

These data demonstrate that eFT226 and hippuristanol exhibit different cellular modes of action that are preserved across cell models (**Fig. 2g**). These findings support a model in which individual mRNAs contain distinct sequence features activating or requiring different functions of eIF4A1 and thus rendering them distinctly sensitive to eIF4A1-inhibtors^1^ (**Fig 2h**).

## Hippuristanol and eFT226 bind to distinct pockets of eIF4A1

To probe the link between the identified mRNA features and compound-inhibited eIF4A1 functions, we needed to understand the structural basis of eFT226 and hippuristanol inhibition in more detail. While a crystal structure of eFT226-related rocaglamide A (rocA) bound to an eIF4A1-AMPPNP-RNA complex has been reported previously^11^, high resolution structural information on hippuristanol binding to eIF4A1 was not available. Here, we report the crystal structure of apo-eIF4A1 C-terminal domain (apo eIF4A1-CTD**, Supplementary Fig. 4a**) and its complex with hippuristanol (**Fig. 3a, Supplementary Fig. 4b-c**) which enabled us to directly compare the structural basis of the mechanism of action of hippuristanol and rocA.

**Figure 3.**
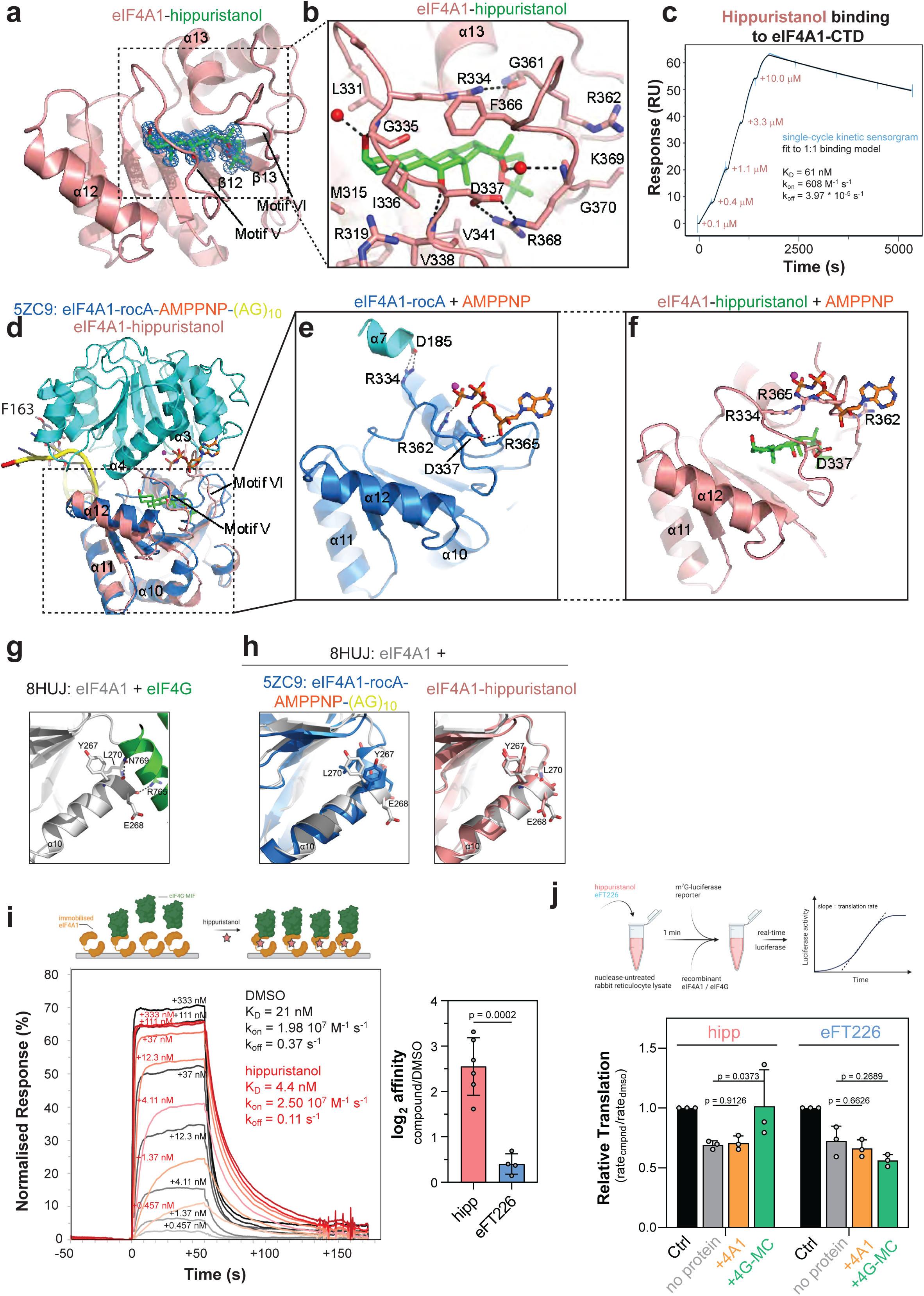
Structure of the eIF4A1-CTD-hippuristanol complex. a, Overall structure of eIF4A1-CTD (salmon) bound to hippuristanol (green, 2|Fo| − |Fc| electron density map in blue and contoured at 1σ). **b**, Close-up view of the binding site highlighting residues within 5Å of hippuristanol.**c**, Representative SPR-sensorgram of hippuristanol binding to immobilised, avi- tagged eIF4A1-CTD. Concentrations given in the plot indicate time points of hippuristanol injections. Six repeat experiments have been performed. **d**, superposition of eIF4A1-CTD- hippuristanol (red) with eIF4A1 in complex with rocA, AMPPNP and an (AG)_10_ RNA (blue, PDBid 5ZC9), and **e-f** close-ups of the hippuristanol binding site in both structures focusing on the conformation of the ATP binding site. **g**, eIF4A1-eIF4G complex (PDBid 8HUJ) showing the primary interface formed by α10 within the eIF4A1-CTD, and highlighting residues involved in hydrogen bonding interactions. **h**, eIF4A1-CTD from the eIF4A1-rocA-AMPPNP-(AG)_10_ (blue, PDBid 5ZC9) and eIF4A1-CTD-hippuristanol complexes (red) superimposed on eIF4A1-CTD from the eIF4A1-eIF4G complex (grey, PDBid 8HUJ) highlighting the positions of residues implicated in hydrogen bonding interactions in the eIF4A1-eIF4G complex. **i**, Representative SPR-sensorgram (normalised to R_max_ = 100%) of indicated concentrations of eIF4G-MIF binding to immobilised, full length avi-tagged eIF4A1 in the absence and presence of 10 µM hippuristanol. Right panel, log2 changes in affinity quantified from the SPR experiments. Data are mean ± sd (n = 6). Three repeat experiments with a technical duplicate each have been performed and are summarised in Table 2. **j**, Rescue of *in vitro* luciferase reporter translation (linear 5’UTR with CAA-repeats^1^) in rabbit reticulocyte lysate with eIF4A1 or eIF4G-MC following inhibition with hippuristanol or eFT226. Data are mean ± sd from n = 3 repeat experiments. P-values calculated from two-way ANOVA.

The X-ray structure of apo eIF4A1-CTD was determined to 2.73 Å resolution (**Supplementary Table 2**) and adopts a RecA-like fold with three copies in the asymmetric unit (RMSD: chain B on A = 1.08 Å; chain C on A = 0.90 Å; chain B on C= 0.48 Å; **Supplementary Fig. 4a**). Subsequently, the structure of eIF4A1-CTD in complex with hippuristanol was determined to 1.70 Å resolution (**Fig. 3a, Supplementary Table 2**). The structure reveals Hippuristanol integrates into eIF4A1-CTD by binding in a cryptic pocket, which is occluded in apo-eIF4A1-CTD and leads to major displacements of residues of DEAD-box RNA helicase motifs V and VI of the RNA and ATP-binding sites (**Fig. 3a, Supplementary Fig. 4b-c**). Specifically, hippuristanol binds above the central β sheet and displaces α12, motif VI between α13 and β13, and the motif V loop with concomitant unwinding of residues D330 to L332 (**Supplementary Fig. 4b**). The binding site is flanked by largely hydrophobic residues (accounting for 21 out of 29 residues within 5 Å of hippuristanol) (**Supplementary Fig. 4c**). The three free hydroxyl groups on hippuristanol form key hydrogen bonding interactions: the A ring hydroxyl hydrogen bonds to the main chain amide of G335, and a water-mediated interaction with the main chain carbonyl of A333; the C ring hydroxyl hydrogen bonds to the main chain amide of V338; the E ring hydroxyl participates in a water-mediated hydrogen bonding interaction with the main chain amide of R368 (**Fig. 3b, Supplementary Fig. 4d**). In contrast, the spiroketal oxygens are structural and do not participate in hydrogen bonding interactions within the pocket. The motif V and VI loops are effectively bridged by 2 key interactions, which engulf the bound hippuristanol: the side chains of D337 and R368 form a salt bridge, and the side chain guanidinium group of R334 hydrogen bonds to the main chain carbonyl of G361 (**Fig. 3b, Supplementary Fig. 4c-d**). In addition, F366 in motif VI in the eIF4A1-CTD-hippuristanol complex forms a cation-π interaction with R344 and adopts a conformation in which it is pointing inwards. In contrast, F366 in apoform eIF4A1-CTD adopts a conformation in which it is pointing outwards (chain B) or is disordered (chains A and C). With this, the binding site of hippuristanol in the crystal structure correlates with previous NMR and mass-spectrometry-based chemical labelling studies^5,12,13^, and reveals the correct orientation and interaction network of hippuristanol within the eIF4A1-CTD. Our structure also explains the elevated hippuristanol-resistance of a previously characterised eIF4A1 variants G370S, and triple mutant eIF4A1^V338I, Q339G, G363T^ through steric clashes and remodelling of the hippuristanol binding site (**Supplementary Fig. 4e**)^2,12^.

The overall binding mode of hippuristanol results in tight caging of hippuristanol within eIF4A1-CTD through residues R334, D337, R365 and R368 which is supported by molecular dynamics simulations suggesting gate-keeping of compound binding and release by these residues (**Fig. 3b**, **Supplementary Fig. 4f**). Here, SPR measurements demonstrate very slow binding and release kinetics of hippuristanol to eIF4A1-CTD with an overall K_D_ of 61 nM (**Fig. 3c**; K_D, eIF4A1-FL_ = 63 nM, **Supplementary Fig. 4g**). This interaction mode is highly specific as 22*S*-epi-hippuristanol (**Fig. 1a**) binds with 1000-fold reduced affinity with an off-rate increased by 10^5^-fold (**Supplementary Fig. 4h**).

In contrast to hippuristanol, the previous crystal structure of a eIF4A1-RocA- AMPPNP-AG_10_-RNA demonstrates that rocaglamides target a bimolecular cavity, including surface residue F163 in the N-terminal domain of eIF4A1 and the RNA (**Fig. 3d**)^11^. In this structure, residues of motif V R362, R365 and D337 in eIF4A1-CTD are involved in AMPPNP binding (**Fig. 3e**). R334 forms a salt bridge with D185 in α7 in the DEAD motif (motif II) in eIF4A1-NTD, bridging both eIF4A1-CTD and -NTD and thus stabilising the typical, catalytically competent, closed conformation of DEAD-box RNA helicases (**Fig. 3e**). Upon hippuristanol binding to eIF4A1-CTD, these motifs and residues are displaced and unable to participate in ATP and RNA binding (**Fig. 3f, Supplementary Movie**). Motif V sterically clashes with the modelled position of AMPPNP in the closed complex (**Fig. 3f**), with the side chain of R334 hydrogen bonding to G361, and D337 participating in a salt bridge interaction with R368. These two interactions clamp together motifs V and VI, and prevent formation of the ATPase-active and RNA-binding competent closed conformation in the hippuristanol-bound state of full-length eIF4A1 (**Fig. 3d, Supplementary Movie**).

In summary, rocaglamides (rocA, eFT226) and hippuristanol bind to distinct domains of eIF4A1 leading to stabilisation or disruption of the ATP and RNA-binding site, respectively, inducing a gain or loss of eIF4A1 function. Through this, eFT226 and hippuristanol specifically enhance and disrupt functions of eIF4A1, respectively, that are consistent with the mRNA sequence features identified in the Ribo-seq.

## Hippuristanol locks inhibited eIF4A1 in eIF4G

Our structural data suggested that binding of rocA and hippuristanol to eIF4A1 affect its ability to transition between open and closed states. Conformational cycling is not only critical for eIF4A1’s enzymatic activity but is also the foundation for its modulation by cofactors^14,15^. Here, the interaction between eIF4G- eIF4A1 is the only one for which a high-resolution structure has been characterised ^16,17^, revealing that eIF4G stabilises eIF4A1 in a half-open conformation inducing exchange of RNA and ATP^14,16,18^. This conformation is achieved through interactions between eIF4G and both eIF4A1-NTD and -CTD, including α10 that forms a key interface with eIF4G (**Fig. 3g)**^16,17^. In the eIF4A1-AMPPNP-RNA-RocA complex (closed state), the C-terminal end of α10 is kinked (pivoted about L266), and is not compatible with forming a favourable interaction with eIF4G (**Fig. 3h, Supplementary Fig. 4i)**. In contrast, α10 is straight in the eIF4A1-CTD-hippuristanol and apo eIF4A1-CTD crystal structures, which allows transitioning between open and half-open states preserving this key interface (**Fig. 3h, Supplementary Fig. 4i-j**). Thus, we hypothesised that if compound binding to eIF4A1 affects conformational cycling, it could also modulate the eIF4A1-eIF4G interaction.

Investigation of the binding of the middle domain of eIF4G (eIF4G-MIF) to immobilised, full-length eIF4A1 by SPR revealed an increase in binding affinity of 5-fold in the presence of hippuristanol (**Fig. 3i**, **Table 2)**. In strong support, we could only determine the binding affinity between eIF4G-MIF and eIF4A1-CTD quantitatively when hippuristanol was present **(Supplementary Fig. 4k, Table 2**). In contrast, the presence of eFT226 alone did not change the affinity between eIF4G-MIF and eIF4A1 (**Fig. 3i**, **Table 2**). These observations are confirmed in an inverse experiment of eIF4A1 binding to immobilised eIF4G-MC (**Supplementary Fig. 4l)**. Strikingly, the presence of hippuristanol did not affect the affinity between eIF4G-MC and hippuristanol-resistant eIF4A1^G370S^ **(Supplementary Fig. 4l-n**). Mechanistically, the kinetic SPR profile indicates broadly similar on rates, with the differences in affinity due to slower dissociation of the eIF4G-eIF4A1 interaction in the presence of hippuristanol (**Fig. 3i**, **Table 2, Supplementary Fig. 4k)**, which agrees with the stabilisation of an eIF4G-binding competent conformation of eIF4A1 by hippuristanol as supported by the crystal structure. This suggested that hippuristanol not only inhibits eIF4A1 but also locks it onto eIF4G, thereby effectively inhibiting the eIF4F complex as opposed to free eIF4A1. Interestingly, cellular concentrations of free eIF4A1 typically exceed eIF4F by more than 10-fold^1,19,20^. Here, hippuristanol targeting specifically eIF4F could explain the discrepancy between the relatively low IC_50_ of hippuristanol and the generally high availability of eIF4A1 in cells. To test this, we performed rescue experiments of mRNA reporter translation with recombinant protein in eFT226- or hippuristanol-treated rabbit reticulocyte lysate (RRL, **Fig. 3j**). Addition of eIF4G-MC, but not eIF4A1, rescued translation of reporter mRNAs in hippuristanol-treated RRL, while eFT226-treated RRL could not be rescued by either protein (**Fig. 3j**).

Altogether, our structural and biochemical data supported the different mode of action of the compounds with hippuristanol specifically inhibiting the activity of the eIF4A1- eIF4G complex.

## Hippuristanol inhibits mRNA loading by eIF4A1

Hippuristanol inhibiting the eIF4G- eIF4A1 complex could follow a mode of action similar to the inhibitory mechanism of dominant-negative mutants of eIF4A1 by perturbing mRNA recruitment into the pre-initiation complex^21^. To examine if hippuristanol selectively targets the eIF4G-eIF4A1 complex in cells, we analysed the mRNAs bound by eIF4G in MCF7 cells using RIP-seq (**Fig. 4a**, **Supplementary Fig. 5a**). This revealed that mRNAs that preferentially associate with eIF4G, are those mRNAs that are most translationally repressed by hippuristanol but not eFT226 (**Fig. 4b, Supplementary Fig. 5b-c**). Strikingly, mRNA features and 5’UTR sequence motifs correlating with increased eIF4G binding closely aligned with the features and nucleotide composition describing hippuristanol-sensitivity (C/CG-rich motifs, **Fig. 4c-d**). This strongly suggests that the mRNAs targeted by hippuristanol are optimised to bind and activate the eIF4G-eIF4A1 and eIF4F complex (**Fig. 4e**). Consequently, inhibition with hippuristanol inactivates eIF4F and its associated functions through stabilisation of the interaction between hippuristanol-inhibited eIF4A1 and eIF4G leading to reduced RNA binding capacity of eIF4F as discussed previously for dominant-negative mutants of eIF4A1^21^ (**Fig. 4e**). Therefore, mRNAs that are already accommodated into the PIC should show a degree of resistance to hippuristanol treatment. To test this, we compared translation of luciferase reporter mRNA in RRL that was treated with hippuristanol before (pre) or after (post) reporter mRNA was added to the reaction. Strikingly, hippuristanol only inhibited reporter translation when RRL was pre-treated, which contrasted eFT226 (**Fig. 4f**, **Supplementary Fig. 5d**). With this the mode of action, hippuristanol was reminiscent of harringtonine, which traps the initiating 80S over the translational start site, suggesting initiation had already taken place and no further rounds of initiation occur after treatment with harringtonine or hippuristanol (**Fig. 4f**). This strongly supports that mRNAs that were already engaged with the translation initiation machinery are not targeted by hippuristanol, supporting our model.

**Figure 4.**
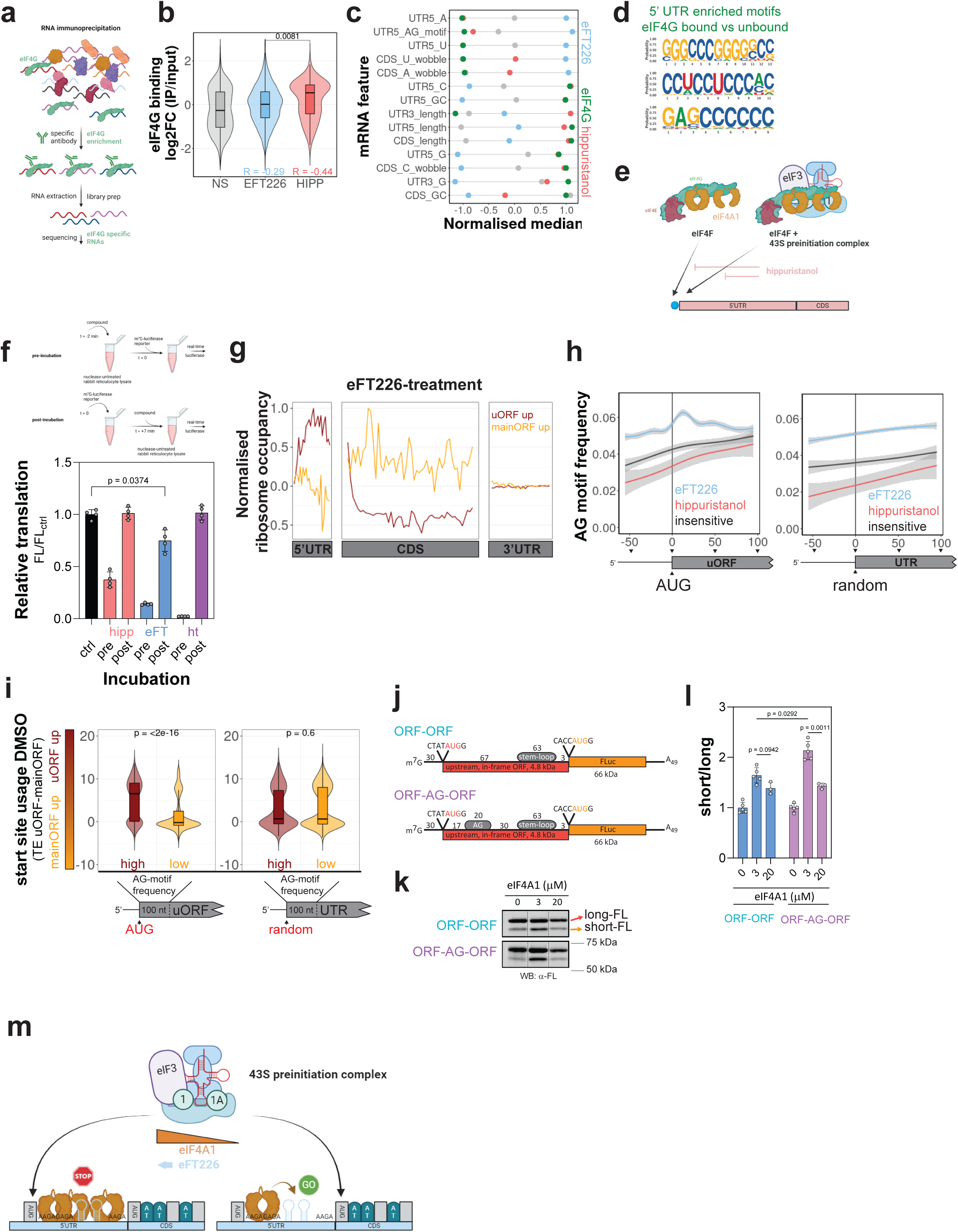
Hippuristanol inhibits loading of mRNAs into eIF4G-eIF4A1 complexes, while eFT226 perturbs translational start-site selection by eIF4A1. a, Schematic presentation of the eIF4G RIP-seq procedure. **b**, Mean log2-fold enrichment in eIF4G-binding (IP / input) of eFT226-, hippuristanol-sensitive and compound-insensitive mRNAs from four repeat experiments. R – Pearson correlation coefficient between eIF4G binding and change in translational efficiency of mRNAs following eFT226 or hippuristanol-treatment. P-value calculated from Wilcoxon-test. Box plots show median ± interquartile range. **c**, Per-feature normalised means of mRNA features with significant difference (p < 0.1) between eFT226- (blue), hippuristanol-sensitive (red) mRNAs, compound-insensitive mRNAs (grey), and of mRNA features linked to eIF4G-binding enrichment (green). **d**, Top three differentially enriched 5’UTR sequence motifs mRNAs enriched in eIF4G-binding against depleted from eIF4G. **e**, Model of the mode of action of hippuristanol. Compound binding to eIF4A1 inactivates the protein, locks it into the eIF4G complex and thus blocks recruitment of mRNA into the 43S pre-initiation complex. **f**, Translation of luciferase mRNA reporter in RRL that is treated with hippuristanol before (pre) or after (post) the mRNA is added to the reaction. Data are mean ± sd from n = 4 repeat experiments. hipp – hippuristanol, eFT – eFT226, ht - harringtonine **g**, Metaplot showing normalised mean counts per million (cpm) of ribosomal occupancy along the 5’UTR, CDS and 3’UTR of mRNAs with increased utilisation of uORF or annotated mainORF translation. **h**, Frequency of AG-motifs downstream of translational starts sites and randomly selected sites in the 5’UTRs of eFT226-, hippuristanol-sensitive and compound-insensitive mRNAs. Data are mean ± 95% ci. **i**, Start site usage in DMSO conditions measured as the difference between the TE of 5’UTR and main-ORFs of mRNAs with high or low AG-motif frequency downstream of start sites or randomly selected positions in the mRNA’s 5’UTR. P-value calculated from Wilcoxon-test. Box plots show median ± interquartile range. **j**, Schematic presentation of the uORF-luciferase mRNA reporters used in panels 4k-l. **k**, Western blot detection of firefly luciferase protein after *in vitro* translation of reporters shown in panel 4j, in rabbit reticulocyte lysate. Full-range blots are shown in Supplementary Figure 5j. **l**, Quantification of western blots representatively shown in panel 4k. Data are mean ± sd from n(Ctrl, 3 µM) = 5, n(20 µM) = 3 repeat experiments. Adjusted p-values for multiple comparison from two-way ANOVA. **m**, Model of eIF4A1 function in alternative start site selection. Depending on the strength and frequence of AG-motifs as well as the eIF4A1 concentration, eIF4A1 unwinds repressive secondary structure to allow translation initiation from the main-ORF, or eIF4A1 clamps onto AG-motifs to block PIC translation and activate uORF utilisation. eFT226 enforces the clamping pathway.

Consequently, mRNA features that promote hippuristanol-sensitivity reflect mRNA properties facilitating selective PIC loading by eIF4A1.

## eFT226 perturbs translational start-site selection by eIF4A1

The reported mode of action of rocaglamides, including eFT226, is to induce sequence-selective clamping of eIF4A1 to AG-rich motifs in the 5’UTR of mRNAs^1,10^. RocA-clamped eIF4A1 blocks translocation of the 43S scanning complex inhibiting initiation from start sites downstream of the clamp-site which in turn favours translation from available open reading frames upstream of the clamp-site (uORF)^10^. The 5’UTRs of eFT226-sensitive mRNAs are enriched for AG-rich motifs (**Fig. 2b** and **2g**) and uORFs (**Supplementary Fig. 5e**), hence, unsurprisingly, eFT226-treatment led to significant changes in start site utilisation (uORF/mainORF) of eFT226-sensitive mRNAs (**Fig. 4g, Supplementary Fig. 5f-g**). Here, our data show that specifically eFT226-sensitive mRNAs, but not hippuristanol-sensitive mRNAs, display positional enrichment of AG-motifs downstream of uORF start sites (**Fig. 4h**), which is not observed at random locations (**Fig. 4h)** nor with other repeat motifs within the same 5’UTRs (**Supplementary Fig. 5h)**. AG-motifs are natural, strong modulators of eIF4A1’s activities related to clamping, multimerisation and unwinding both *in vitro* and in cells^1,22^ and their occurrence is increased in uORF-containing 5’UTRs (**Supplementary Fig. 5i**). Expectedly, we find that this specific architecture of start sites and AG-motifs directly impacts start site utilisation also in untreated conditions (**Fig. 4i**) supporting a possible native role of eIF4A1 in this process. This agrees with previous observations that rocA-sensitive mRNAs naturally utilise alternative start site more frequently^10^ and suggests furthermore that it is the location of AG motifs that guides a function of eIF4A1 in influencing translational start site selection. To test this directly, we performed *in vitro* translation assays using mRNA constructs with two alternative translational start sites separated by a stem-loop and AG motif in an otherwise linear CAA 5’UTR (**Fig. 4j**). The results demonstrate a change in translational start site utilisation in an eIF4A1-dose dependent manner, that is enhanced by the presence of an AG-motif (**Fig. 4k-l**) and strongly stimulated in the presence of eFT226, while hippuristanol inhibits reporter translation globally (**Supplementary Figure 5j-k**).

Together, this supports a role of eIF4A1 in translational start site selection of mRNAs harbouring features enriched in eFT226-sensitive mRNAs utilising unwinding and clamping functions induced by AG motifs (**Fig. 4m**).

## Validation of the mRNA sequence-function relationship of eIF4A1

To test and validate our model of the mRNA sequence – eIF4A1 function relationship (**Fig. 4e, 4m**) , we performed a massively parallel reporter assay i.e., Nascent-Peptide Translating Ribosome Affinity Purification assay (NaP-TRAP)^23^, to measure reporter mRNA translation in MCF7 cells. Briefly, cells were transfected with an mRNA reporter library encoding a FLAG-GFP fusion protein. The translation efficiency of individual reporters was assessed by measuring reporter-enrichment following pulldown of the FLAG-tag, normalized to input levels using high throughput sequencing (**Fig. 5a**). In this set up, the enrichment of each reporter correlates with the number of ribosomes actively translating the FLAG-tag-containing ORF, thereby reflecting its translation efficiency. To specifically investigate the impact of 5’UTR sequence features, all mRNA reporters in the library were designed with identical CDS and 3’UTR sequences, as well as consistent 5’UTR- and total mRNA-lengths. Consequently, the only variable across reporters was the 5’UTR sequence (**Fig. 5a**). Each 5’UTR contained a 124-nt variable region derived from zebrafish 5’UTRs, resulting in a library of 11,088 unique 5’UTR sequences.^23^ This approach allowed us to not only test our model but also examine if the modes of actions of eFT226 and hippuristanol are faithful across mRNA sequences from a different evolutionary origin.

**Figure 5.**
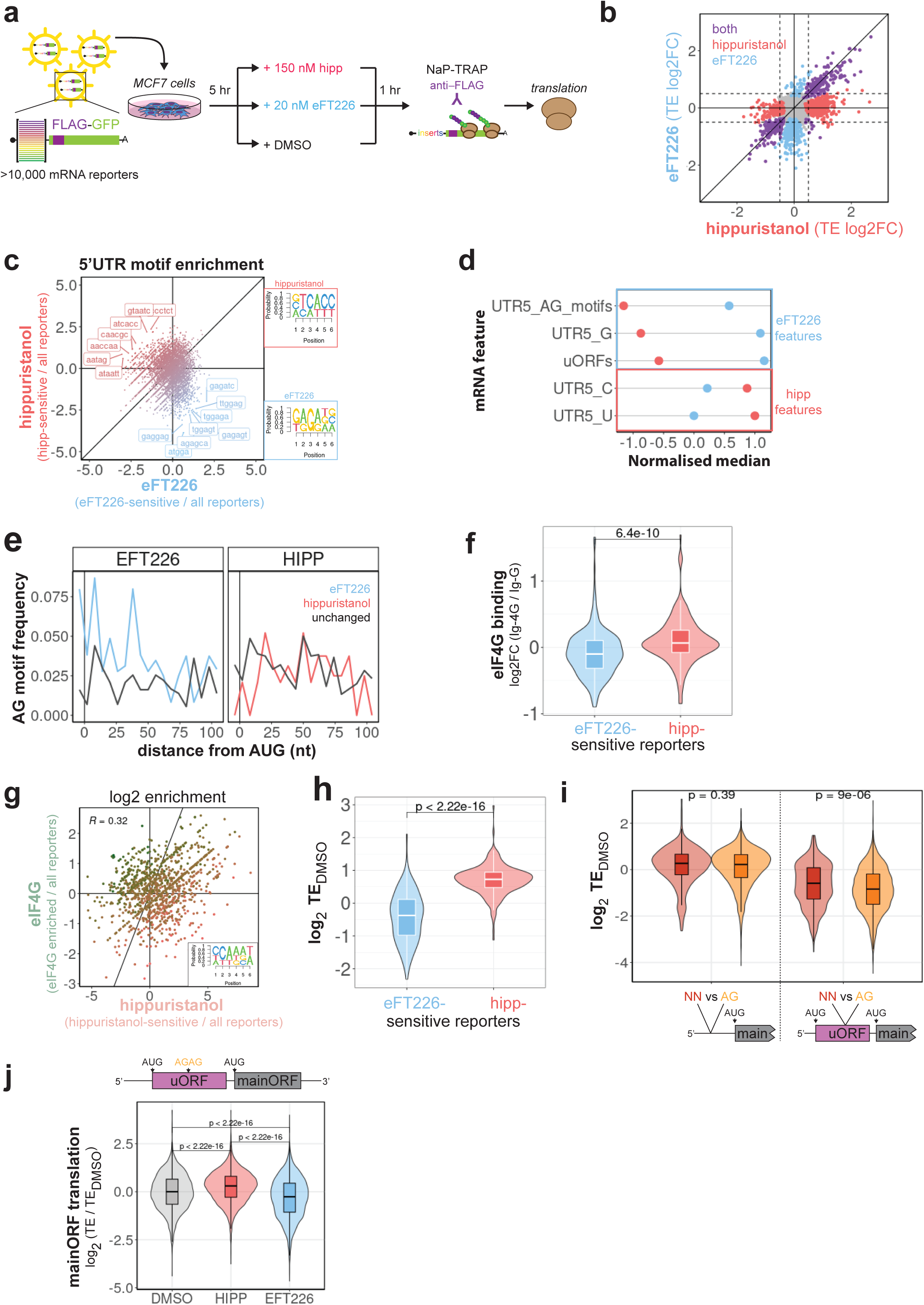
**Validation of the mRNA sequence – eIF4A1 function relationship**. **a**, Schematic presentation of the NaP-TRAP assay. **b,** Mean log2-fold changes in translational efficiencies (TE) of reporters following eFT226- and hippuristanol-treatment. Data from four repeat experiments are shown. **c**, Log2-enrichment (compound-sensitive reporters / all reporters) of 5-6mer sequence motifs within the 5’UTRs of hippuristanol and eFT226-sensitive reporters. Top differentially enriched motifs are labelled. the insert shows the consensus sequence of the top three enriched 6mer-motifs. **d**, Per-feature normalised medians of mRNA features with significant difference (p < 0.1) between eFT226- (blue) and hippuristanol-sensitive (red) reporters. **e**, Frequency of AG-motifs downstream of AUG starts sites in the 5’UTRs of eFT226- and hippuristanol-sensitive and compound-insensitive reporters. **f**, Mean log2-fold enrichment in eIF4G-binding (eIF4G-IP / IgG-IP) of eFT226-, hippuristanol-sensitive mRNAs reporters. n = 4. P-value calculated from Wilcoxon-test. Box plots show median ± interquartile range. **g**, Log2-enrichment (hippuristanol-sensitive or eIF4G-enriched reporters / all reporters) of 4-6mer sequence motifs within the 5’UTRs of hippuristanol-sensitive reporters and reporters enriched for eIF4G binding. The insert shows the consensus sequence of the top five co- enriched 6mer-motifs. R – Pearson correlation coefficient. **h**, Mean translational efficiencies in DMSO control conditions of hippuristanol and eFT226-sensitive reporters. P-value calculated from Wilcoxon-test. Box plots show median ± interquartile range. **i**, Effect of the presence of 5’UTR AG-motifs on the translational efficiencies of reporters in DMSO control conditions with or without uORFs (scheme) implicating differential uORF activation. P-value calculated from Wilcoxon-test. Box plots show median ± interquartile range. **j**, log2-difference in main ORF translation of reporters with AG-motifs downstream of uORF within their 5’UTR (scheme) under hippuristanol or eFT226-treatment relative to DMSO control. Box plots show median ± interquartile range.

MCF7 cells were first transfected with the reporter library and translation allowed to proceed for 5 h before cells were treated with DMSO, hippuristanol or eFT226 for one hour before harvest and NaP-TRAP library preparation and sequencing (**Supplementary Fig. 6a- b**). Following an analogous analysis strategy as with the Ribo-seq data, we identified pools of mRNAs sensitive to hippuristanol or eFT226 only (**Fig. 5b**), that were characterised by a set of molecular 5’UTR features very reminiscent of their respective group in the Ribo-seq data: The 5’UTRs of eFT226- or hippuristanol-sensitive reporters were differentially enriched for 5-6mer RNA sequence motifs (**Fig. 5c**), with eFT226-sensitive 5’UTRs showing specific enrichment for AG-motifs (**Fig. 5c-d, Supplementary Fig. 6c**), while hippuristanol-sensitive 5’UTRs were enriched in Cs (**Fig. 5c-d, Supplementary Fig. 6d**). Moreover, eFT226- sensitive 5’UTRs contained increased numbers of alternative start sites (**Fig. 5d**, **Supplementary Fig. 6e**) with positional enrichment of AG-motifs downstream of those sites (**Fig. 5e, Supplementary Fig. 6f)** that was absent in hippuristanol-sensitive 5’UTRs **(Fig. 5e)**. eIF4G-RIP-seq of the reporter library (**Supplementary Fig. 6g-h)** revealed that association between reporters and eIF4G correlated with their sensitivity to hippuristanol- but not eFT226-treatment (**Fig. 5f, Supplementary Fig. 6i,** consistent with **Fig. 4b**).

Additionally, 5’UTR motifs associated with (enriched) eIF4G binding were also found to be enriched in hippuristanol- (**Fig. 5g)** but not eFT226-inhibited reporters (**Supplementary Fig. 6j).** As a result of these individual mRNA features, hippuristanol-sensitive reporters showed a generally higher translational efficiency together with their increased eIF4G binding (**Fig. 5h**), while the presence of AG-motifs only repressed (main ORF) translation specifically in combination with alternative start sites (**Fig. 5i**, consistent with **Fig. 4i**). In support of our model, the latter effect is limited after loss-of-function of eIF4A1 following hippuristanol- treatment (**Fig. 5j**) and, conversely, enhanced by gain-of-function after eFT226-treatment (**Fig. 5j**). Thus, 5’UTR features enriched in hippuristanol- and eFT226-sensitive reporters agree with eIF4A1 functions in mRNA-loading and start site-selection, respectively, as derived from our Ribo-seq data supporting our model.

Concluding, we find a consistent relationship between mRNA sequence and eIF4A1 function across mRNA sequences from distinct evolutionary origin.

## Evolutionary anchorage of the mRNA sequence-eIF4A1 function relationship

The results from the NaP-TRAP assay showed a consistent relationship between mRNA sequence and eIF4A1 function across mRNA sequences from zebrafish and human. This prompted us to examine if our model shows evolutionary conservation. eIF4A is an highly conserved protein with 71% sequence identity across the phylogenetic tree (**Fig. 6a**) and nearly identical, AlphaFold2-predicted structures with average RMSD between structures smaller than 0.5 Å (**Fig. 6b** and **Supplementary Fig. 7a**). Moreover, eFT226- and hippuristanol-binding sites are conserved, except for *C. elegans* which contains a glycine at the position homologue to F163 that binds eFT226 (**Supplementary Fig. 7b**). Based on the general high degree of conservation, eIF4A is considered to participate in translation initiation performing conserved functions^24^. However, there are remarkable biochemical differences between for example *S. cerevisiae TIF1* and human *EIF4A1*^24^, the functional consequences of which are incompletely understood.

**Figure 6.**
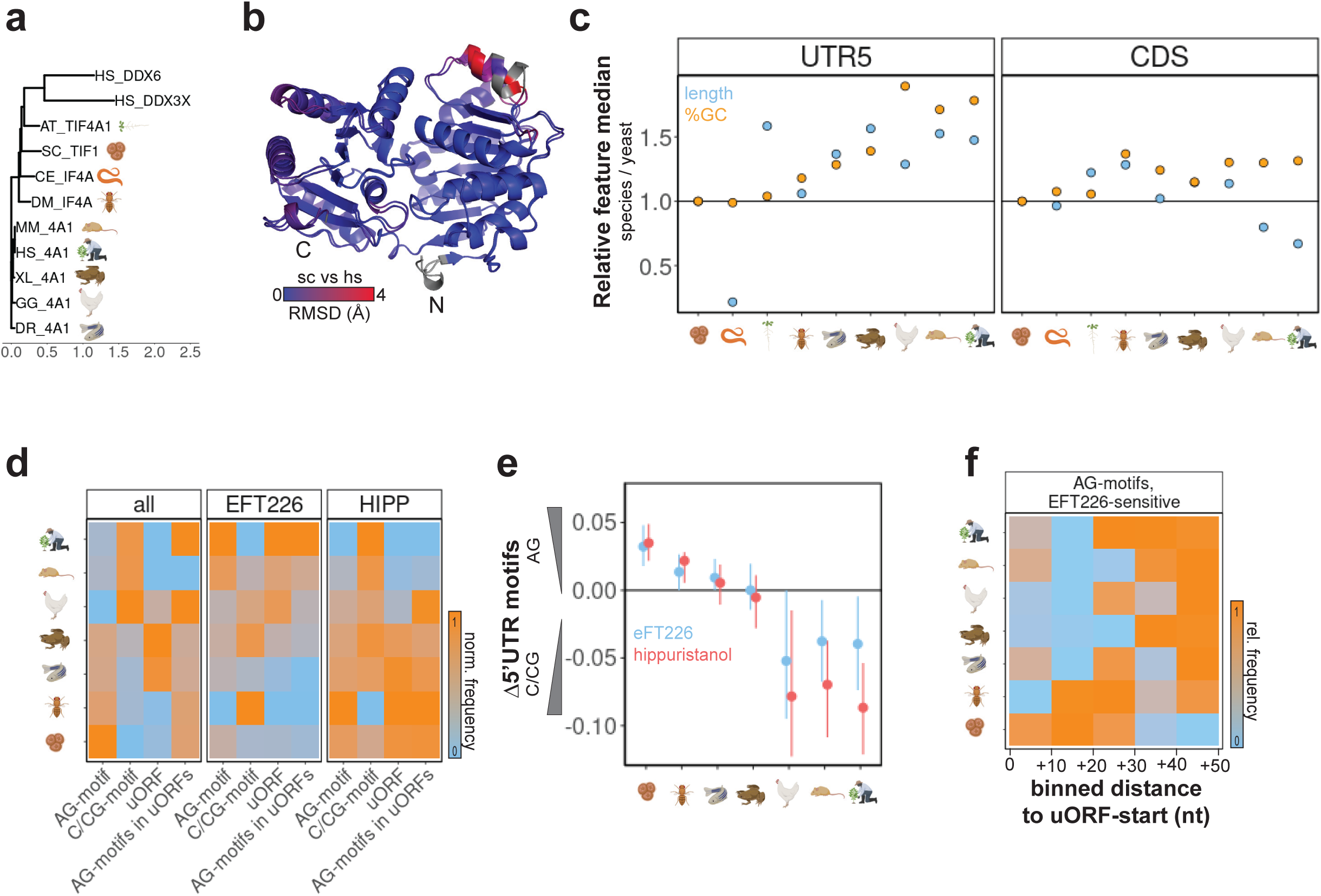
**Evolution of the mRNA sequence – eIF4A1 function relationship**. **a**, Phylogenetic tree based on the primary sequence of eIF4A proteins from *Gallus gallus* (GG), *Mus musculus* (MM), *Homo sapiens* (HS), *Danio rerio* (DR), *Drosophila melanogaster* (DM), *Arabidopsis thaliana* (AT), *Xenopus leavis (*XL), *Caenorhabditis elegans* (CE) and *Saccharomyces cerevisiae (*SE). Human DDX3X and DDX6 are included for reference. **b**, AlphaFold2-predicted structures of *S. cerevisiae* (sc) and *Homo sapiens* (hs) eIF4A coloured by the root mean square difference (RMSD in Å) between in the structures. **c**, Median of length and GC-content of 5’UTRs and CDS regions relative to the feature median of *S. cerevisiae*. **d**, Frequency of indicated 5’UTR features normalised within each feature and group. Grouping based on annotation of human-orthologue transcripts. **e**, Median of the difference in the frequency between AG- and C/CG-motifs in the 5’UTRs of eFT226- and hippuristanol- sensitive transcripts (homology based) of indicated species. **f**, Frequency of AG-motifs in 10- nt bins downstream of alternative start sites in 5’UTRs of indicated species normalised within each species.

Given that 5’UTRs show more variation in sequence composition and length than coding regions across evolution (**Fig. 6c**)^25^, it is not very surprising that the general level of positional conservation of 5’UTRs of the different species relative to human is substantially poorer than conservation between coding regions (**Supplementary Fig. 7c**). While the level of positional conservation is also not different between eIF4A1-dependent or eIF4A1- indepedent 5’UTRs (**Supplementary Fig. 7d**), it gradually improves over time, indicating gain and potential selection of sequence features. Indeed, focusing the analysis on mRNA features that we linked to specific eIF4A1 functions i.e., C/CG-motif-, AG-motif and start site frequency, we observe evolutionary trends (**Fig. 6d, left panel**). While the overall occurrence of AG-motifs decreases along the evolutionary time scale (**Fig. 6d, left panel**), it increases specifically downstream of start sites, with the frequency of alternative start sites being maintained (**Fig. 6d, left panel**). Strikingly, these features become particularly enriched within eFT226-sensitive mRNAs (**Fig. 6d, right panels**), while hippuristanol- sensitive mRNAs follow the opposite trend and gain CG-rich motifs together with losing alternative start sites (**Fig. 6d, right panels**). Here, our analysis shows that eFT226- homologue transcripts specifically maintained a higher frequency of AG-motifs than hippuristanol-homologues (**Fig. 6e**), which suggests co-development of basic mechanisms of eIF4A1 function alongside the mRNA sequences. It has been previously discussed for rocA-induced (eFT226-derivative), eIF4A1-mediated utilisation of alternative start site, that optimal positioning of these AG-motifs occurs at approximately 18 nt downstream of start sites^10^. This would allow better identification and activation of alternative start sites within the pre-initiation complex A-site. Interestingly, we observe that the distance between alternative start sites and AG-motifs within the 5’UTR of eFT226-senstive mRNA-homologues increases over evolution to a location of approximately +20 nt (**Fig. 6f)**. This is in line with the observed, decreasing content of AG-motif across evolution (**Fig. 6d** and **Supplementary Fig. 7e)** and further indicates a tuning of eIF4A1 functions relative to the genomic background, that increases in length, and changes in nucleotide composition over time (**Fig. 6c**). The overall evolutionary trend is confirmed by unbiased and independent clustering of the data (**Supplementary Fig. 7f**) and suggests that functions of eIF4A have evolved in accordance with selection for and gain of information in the genomic background of the 5’UTRs.

## Discussion

eIF4A1-dependent mRNAs have been investigated previously in different studies using a variety of cell lines and various strategies including siRNA-mediated knockdown, genetic interference or chemical inhibition of eIF4A1^2,6–8,10,26^. While each study identifies mRNA sequence features involved in eIF4A1 activity, data sets and features correlate only poorly and how they relate to specific eIF4A1 functions is not investigated^2,6–8,10,26–28^, leaving the question of how eIF4A1 functions to deliver specific translational programmes unanswered. Through in-depth translational profiling of eIF4A1 using two therapeutically important eIF4A1-inhibitors, we have not only characterised the inhibitors’ biochemical and cellular mode of action but also uncovered the mechanisms by which mRNA sequences are linked to eIF4A1 function. While each of the inhibitor-sensitive mRNAs are *per se* eIF4A1- dependent (**Fig. 1**), we identified at least two groups within those mRNAs harbouring distinct sequence features that activate specific eIF4A1 functions (**Fig. 2**). This suggests that previous studies may have characterised different eIF4A1-dependent subgroups in a condition-dependent manner which may contribute to apparent discrepancies across these studies^2,6-8,10,26-28^.

Our presented work links mRNA sequences with functions of eIF4A1 in mRNA loading and alternative start site selection. Specifically, we provide evidence that 5’UTR AG- motifs participate through a context-dependent dual role in alternative start site selection by eIF4A1. Depending on the strength/frequency of the AG-motif in recruiting eIF4A1, the AG- motifs either activate eIF4A1 unwinding of repressive RNA structure with subsequent PIC translocation and initiation at the downstream ORF(s) (**Fig. 4m**)^1^, or induce clamping of eIF4A1 thereby blocking PIC translocation and facilitating initiation from upstream ORF(s)^10^. This model is strongly supported by 1) a recent study describing that eIF4A1 appears to be involved in bi-directional scanning of the PIC which affects stringency of start site selection, however, the mechanisms relating to eIF4A1 function were not investigated^29^, and 2) a recent cryo-EM structure identifying an eIF4F-independent eIF4A1 molecule clamped to an AG-motif at the entry-site of the mRNA channel of the preinitiation complex^30^. Here, our data indicates an optimal localisation of AG-motifs and clamped eIF4A1 just ∼ 15-20 nt downstream of the P-site of the scanning PIC (**Fig. 4g** and **6f**), which is exactly the distance of the eIF4A1 molecule from the AUG-containing P-site of the PIC in the cryo-EM structure^30^ and could provide a mechanism for increased alternative start site utilisation similar to what has been described for rocA-induced uORF activation previously^10^.

In our study, we provide the first high-resolution structure of hippuristanol bound to the C-terminal domain of eIF4A1 revealing atomic details of the distinct modes of action of hippuristanol in comparison to eFT226 (**Fig. 3a**). The structure highlights that hippuristanol binds to a cryptic pocket in the C-terminal domain of eIF4A1 and is gatekept by a series of arginine residues that open and close the binding pocket. This is supported by the slow kinetics of the binding process. Consequently, hippuristanol binding displaces loops critical for ATP- and RNA-binding, which supports previous biochemical observations of loss-of-function of eIF4A1 by compound activity^2,5^. In contrast, rocaglamides, including eFT226, target a bimolecular cavity specifically formed by eIF4A1-NTD and a sharply bent pair of consecutive purines in the RNA, supporting the unique gain-of-function mode of action of these compounds^11^. In addition, our structure highlights a unique and undiscovered feature of the mode of action of hippuristanol by which compound-binding to eIF4A1 stabilises the interaction between eIF4A1 and eIF4G directly linking hippuristanol-induced translational inhibition to eIF4A1’s mRNA loading function within the eIF4F complex. Consequently, the mode of action of hippuristanol involves an unprecedented specificity to induce loss-of-function of the eIF4F complex. To date, rocaglamides have been the class-of-choice for chemical evolution of eIF4A1-inhibitors due to the lack of high-resolution, structural information on other classes of compounds. Our structure agrees with previous studies that mapped the hippuristanol-binding site within the C-terminal domain of eIF4A1^12,13^ and can inform the development of hippuristanol analogues with improved physiochemical properties via structure-based design^31^.

Our model of the mRNA sequence – eIF4A1 function relationship is based on a comprehensive set of experiments integrating data from human and mouse cells, and their careful interpretation. These two systems were chosen to increase confidence in the analytical output through more depth and variance of the input sequence information as their homologue transcripts show a reasonable degree of divergence, specifically in their 5’UTR (**Fig. 6c and Supplementary Fig. 7b**). The correlations of the different data sets in this presented study varies in the range of 0.4 - 0.6 indicating that certain aspects of the model could be over- or underestimated. Here, we note that due to time-restrictions in the TMT-pSILAC, this data set is only representative of the topmost translated proteins, which could contribute to the less-than-expected correlation with our other data. Nevertheless, mRNA feature analysis based on this data set extracts the identical features as found with the Ribo-seq data. Further, we do not exclude that hippuristanol does not target steps in translation initiation downstream of mRNA recruitment to the PIC, or that eFT226 does not have an indirect effect on eIF4F activity itself, as suggested previously for chemically related drugs^5,27^. As our data are consistent between experimentally tested human and mouse cells, we consider our model to represent the main effect directly incurred by the compound activity on eIF4A1.

We addressed validation of the mRNA sequence – eIF4A1 function relationship through a massively parallel reporter assay in which mRNA reporters contained a 124 nt variable region within their 5’UTR with identical cap-proximal nucleotides as well as coding and 3’UTR regions. With this, differences in reporter translation were directly correlated to steps in translation initiation. The results from this assay replicated and thus support all key points of our model: i) distinct eFT226- and hippuristanol-sensitive reporters with unique sequence features (compare **Figures 2g** and **5b-d**), ii) hippuristanol- but not eFT226- inhibition of reporters correlated with the capacity of reporters binding to eIF4G (compare **Figures 4b** and **5f**), the frequency of 5’UTR AG-motifs controls utilisation of alternative start sites in an eIF4A1-dependent manner (compare **Figures 4h-i** and **5e,j**). In further support, hippuristanol-sensitive reporters showed increased translational efficiency over eFT226- sensitive mRNA reporters, which is directly attributed to increased reporter recruitment to eIF4G and thus PICs despite identical 5’ ends and overall 5’UTR length between the reporter 5’UTRs (**Fig. 5h**). However, this could explain the poorer correlation between hippuristanol-sensitivity and eIF4G-binding observed for the reporters as compared to the endogenous mRNAs (compare **Supplementary Figures 5b-c and 6i**). Moreover, while 5’UTR sequence motifs related to this mRNA loading activity both from the reporter assay and Ribo-seq matched their paired eIF4G-RIP-seq, the discovered motifs themselves were slightly divergent in nucleotide compositions between the assays (6-mer consensus sequences *d. rerio*: CCAAAT, **Fig. 5g**; *h. sapiens*: CCCCCG **Fig**. **4d**). For this reporter assay, the nucleotide sequences used to build the 5’UTR library were designed to contain 5’ leader information unrelated to human origin. In our case, we chose the zebrafish transcriptome as its genomic background is distinct from human^23^. Hence, we hypothesised that a species-specific genomic background explains the observed differences between the Ribo-seq and reporter assay. In agreement, we observe an evolutionary trend that 5’UTRs used to be A(G)-rich in yeast and gained more C(G)-content (**Fig. 6d**). Interestingly, it is specifically eFT226-homologue mRNAs that accumulate alternative start sites and maintain a high frequency of AG-motifs downstream of these start sites (**Fig. 6d-e**). In contrast, hippuristanol-homologue mRNAs become depleted for these features and gain or maintain sequence compositions possibly lining up with their ability to bind eIF4G. With this our data not only shows (co)evolutionary development of eIF4A functions and its target sequences, but also the preservation of the compound mode of action on eIF4A despite the origin of the reporter sequences. It remains unclear, what pressure drove this evolutionary selection and which biological role it is serving. As we find that these distinct mRNA features are also present in mRNAs coding for functionally related proteins and protein families, we speculate that distinct post-transcriptional, temporal regulation schemes and control over proteo- isoforms could play a role.

eIF4A1 has long been believed to function in an RNA sequence-unspecific manner^32,33^. However, while it was understood that not all cellular mRNAs require eIF4A1 activities and functions to the same extent^1,34,35^, the underlying mechanism of how eIF4A1 functions contribute specifically to establish gene expression programmes remained unclear.

Our previous and current works provide insight towards resolving this debate by demonstrating how mRNAs are tuned to meet their eIF4A1-requirements to facilitate translation by harbouring regulatory 5’UTR sequence motifs that activate specific eIF4A1 functions^1^. This mRNA sequence – eIF4A1 function relationship is preserved in a different setting such as in the investigated colorectal cancer model (Apc^-/-^, Kras^G12D/+^, Fig 1). In this model, the expression of hippuristanol-sensitive mRNAs is specifically increased over eFT226-sensitive mRNAs (**Supplementary Fig. 3h**), directly suggesting that eIF4A1 functions may be differentially critical to establish proliferative gene expression programmes. As hippuristanol inhibits the mRNA loading function of eIF4A1, we speculate that strongly proliferative mRNAs become translationally upregulated by disease-specific increase in ribosomal recruitment to these mRNAs.

Concluding, our model of the mRNA sequence – eIF4A1 function relationship not only illustrates how RNA helicase activity is deployed mRNA specifically but also adds a new layer to mRNA self-regulation and orchestration of gene expression programmes through matching the activities of translation factors with presence of mRNA sequence motifs. With this our data also delivers guidelines to sequence-specifically optimise translational efficiencies of mRNAs that could be exploited to increase productivity of antigens from therapeutic mRNAs.

## Material and Methods

### Reagents

Enzymes: TEV protease (Thermo Fisher Scientific 12575015) RNase I (cloned, 100 U/µL, Thermo Fisher Scientific, AM2295), T4 PNK (NEB M0201S), Benzonase (Merck E1014),DNaseI (Merck DN25-10MG), 2X KAPA HiFi HotStart ReadyMix (Roche #7958935001), SuperScript™ III Reverse Transcriptase (Thermo # 18080044), RNase H (NEB # M0297S), RNase I (NEB # M0243S), DNase I (NEB #M0303L), RNaseOUT (Thermo #10777019) Antibodies: eIF4A1 (ab31217, Abcam UK), puromycin (Merck MABE343), eIF4G (for blotting Cell Signalling Technologies 2498S, RIP (Life Technologies MA5-14971), PABP (abcam ab21060), eIF4E (BD biosciences 610270), actin (Fisher Scientific 11373069), firefly luciferase (Proteintech 27986-1-AP), IgG-rabbit (Cell Signalling Technologies 2729S) Kits: HiScribe™ T7 mRNA Kit with CleanCap® Reagent AG (NEB E2080S), Rabbit Reticulocyte Lysate (Promega L4151), TMT 16plex reagent kit (A44522 Thermo Scientific). Nextflex small RNA v3 (Revvity, NOVA-5132-06), RiboCop rRNA Depletion Kit HMR V2 (Lexogen 144.96), Corall Total RNA-Seq Library Prep Kit (Lexogen 095.96), λDE3 Lysogenization Kit (Merck 69734-3), mMESSAGE mMACHINE™ SP6 Transcription Kit (Thermo #AM1340), RNA Clean & Concentrator-5 (Zymo #R1013), Zymoclean Gel DNA Recovery Kit (Zymo #D4001), DNA Clean & Concentrator-5 (Zymo #D4004), Qubit™ 1X dsDNA High Sensitivity (Thermo # Q33230) RNAs: all RNAs were purchased from IDT. Columns for FPLC: HisTrap HP 5 mL (17524802, Cytiva), HisTrap FF 5 mL (17525501, Cytiva), Superdex 75pg 26/60 (28989334, Cytiva), Superdex 200pg 26/60 (28989336, Cytiva). eIF4A-inhibitors: hippuristanol (gift from John Le Quesne), hippuristanol used in structural studies and SPR was purchased from Pharmaron, eFT226 (HY-112163, Generon UK). SILAC amino acids: Lys-^12^C ^14^N (Lys0), Arg-^12^C ^14^N (Arg0), Lys-^13^C ^15^N (Lys8), Arg-^13^C ^15^N (Arg10) all purchased from Cambridge Isotope Laboratories (#ULM-8766, #ULM-8347, #CNLM-291-H, #CNLM-539-H).

Chemicals: cOmplete, EDTA-free Protease Inhibitor Cocktail (Roche 11873580001), n-Dodecyl-β Maltoside (Thermo Fisher 89903), SIGMAFAST™ EDTA free protease inhibitor cocktail (Merck S8830), puromycin (Invivogen ant-pr-1), Phenylmethanesulfonyl fluoride (Merck P7626-25G), Lipofectamine Messenger Max (Thermo #LMRNA003), Anti-FLAG magnetic beads (Thermo # A36797), TRIzol (Thermo #10296010), GlycoBlue (Thermo #AM9516), Chloroform (Sigma #CX1060), Isopropanol (American Bio # AB07015-01000), AMPure XP (Beckman Coulter #A63880), DNPTs (NEB #), SYBR® Green I nucleic acid gel stain ( Sigma #S9430), D1000 ScreenTape (Agilent #5067-5582), D1000 Reagents (Agilent #5067-5583)

### Cell lines and auth

MCF7 cells were purchased from ATCC and were already authenticated. MCF7 cells were additionally authenticated by Eurofins using PCR-single- locus-technology. All cell lines were regularly tested for mycoplasma. All tests were negative and confirmed the absence of mycoplasma contamination.

### Organoid isolation and culturing

Experiments were performed under license from the UK Home Office (PP3908577). *VillinCre*^ER^ *Apc*^fl/fl^ *Kras*^G12D/+^ (referred to as Apc^-/-^KRAS^G12D/+^) mice were generated (all colonies inbred C57BL/6J, generation >= 7, induced at 8 - 15 weeks), organoids isolated (lines 240614 - female, 240615 - female, RGH60.1g - female for eFT226, RVJ22.5c - male, RVJ22.5h - female, RVJ22.6d - female for hippuristanol experiments) and cultured as described previously^36^. Sex was not considered in the analyses. Ribosome profiling in organoids was performed in three different organoids lines per inhibitor derived from different animals.

### Biological Resources

bacterial strains: *E. coli* BL21 (DE3) CodonPlus-RP (#230255, Agilent), *E. coli* BL21 (DE3) CodonPlus-RIPL (#230280, Agilent), *E. coli* CVB101 (#CVB101, Avidity). Plasmids: pGL3 (Promega). Cell lines: MCF7 (HTB-22, ATCC) **Ribooligonucleotides**. RNAs used in this study were purchased from IBA Lifescience and Integrated DNA Technology and are listed in **Supplementary Table 1**.

### Purification of recombinant proteins for *in vitro* assays

Protein production and purification was performed as described previously^1^. eIF4G-MIF corresponds to residues 712-1011 and eIF4G-MC to residues 674–1600.

### RNA binding and release

Experiments were performed essentially as described previously^1^. eIF4A1 was pre-incubated with 20 µM inhibitors in reaction buffer in the absence of ATP for 30 min. RNA-binding was then initiated by the addition of ATP. For RNA release experiments, complexes were pre-formed between 2 µM eIF4A1 and 50 nM RNA in assay buffer in the presence of ATP before ATPase-dependent RNA release was monitored in the presence or absence of 20 µM inhibitors.

### RNA unwinding and EMSA

Fluorescence-based RNA unwinding and EMSA experiments were performed as described previously^1^ in the presence of 20 µM inhibitors.

### IC50 determination

MCF7 cells were seeded at a density of 25000 cells per 96-well in DMEM (Gibco) supplemented with 10 % FBS (Gibco) and 2 mM final concentration of L-glutamine (Gibco) in for 24 h. Medium was replaced with fresh medium containing indicated concentrations of eIF4A1-inhibitors at a final concentration of 0.1% DMSO in technical duplicates (two wells per concentration). For titration experiments with organoids, one 96- well with one plug of organoid was used per concentration. After 1 h treatment, puromycin (Invivogen) was added to the medium at a concentration of 10 µg/mL. After 10 min incubation cells were washed with PBS and lysed in IC-buffer (150 mM sodium chloride, 1.0% NP-40, 0.5% sodium deoxycholate, 0.1% SDS, 50 mM Tris, pH 8.0) in the presence of protease inhibitors (Roche 11873580001). Lysates were cleared by centrifugation and transferred onto a nitrocellulose membrane pre-equilibrated in IC-buffer using a dot plot apparatus (BioRad). After transfer, the membrane was blocked in 7% milk and puromycin incorporation detected by incubation with anti-puromycin antibody for 2 h, followed by incubation with a secondary anti-mouse antibody for 1 h and subsequent scanning of the membrane using an Odyssey Imager (Licor). Signals were quantified using ImageStudio Lite v5.2.5 software (Licor).

### Reporter mRNA construction and *in vitro* transcription

mRNA reporters were constructed as reported previously^1^.

### *In vitro* translation assays

Experiments were performed as described previously or, when uORF reporters were used, with minor modifications^1^. In the latter case, translation reactions were stopped by addition of SDS-gel loading buffer and volume equivalent to 1 uL of the reaction loaded on 10% SDS-polyacrylamid gels. Proteins were transferred to nitrocellulose membranes by western blotting. Produced firefly luciferase was detected via fluorescence using Odyssey Licor after incubating the membranes with anti-firefly (Proteintech) and anti-rabbit (CST) antibodies. Detected bands were quantified using ImageStudio Lite v5.2.5 (Licor).

### Protein expression and purification for eIF4A1-CTD crystallography

Human eIF4A1 C-terminal domain (residues 239-406; eIF4A1-CTD) was cloned into the pETNHT vector and expressed as a TEV cleavable N-terminal His_6_-tag fusion protein in BL21-CodonPlus (DE3)- RIPL *E coli* cells. Cell cultures grown in LB medium, supplemented with ampicillin, were induced with 0.5 mM IPTG overnight at 18 °C. The cells were harvested by centrifugation and the pellets washed with ice cold PBS before storing at -20 °C.

Cells were resuspended in Lysis buffer (20 mM Tris-HCl pH 7.5, 500 mM NaCl, 5 % glycerol, 1 mM DTT, SIGMAFAST™ EDTA free protease inhibitor cocktail, Benzonase® nuclease) and lysed using a Constant Systems cell disruptor. The lysate was clarified by centrifugation, and the filtered supernatant was applied to a HisTrap HP column equilibrated with IMAC buffer (20 mM Tris-HCl pH 7.5, 500 mM NaCl, 5 % glycerol) supplemented with 20 mM imidazole. The column was washed sequentially with IMAC buffer supplemented with 20 mM imidazole, High salt buffer (20 mM Tris-HCl pH 7.5, 1 M NaCl, 5 % glycerol), IMAC buffer supplemented with 20 mM imidazole and IMAC buffer supplemented with 40 mM imidazole. The protein was eluted with a linear gradient of 40-500 mM imidazole in IMAC buffer.

The eluted protein was diluted 1:1 v/v with Dialysis buffer (20 mM Tris-HCl pH 7.5, 300 mM NaCl, 5 % Glycerol, 1 mM DTT), treated with TEV protease and, using a 10 kDa molecular weight cut off tubing, dialysed against Dialysis buffer overnight at 4 °C. The dialysed TEV-treated protein was reloaded onto a HisTrap HP column equilibrated with IMAC buffer. The untagged eIF4A1-CTD protein was collected in the flow-through and wash fractions, concentrated and applied to a HiLoad 26/60 Superdex 75 pg size exclusion chromatography column equilibrated with 20 mM Tris-HCl pH 7.5, 300 mM NaCl, 5 % Glycerol, 1 mM DTT. The purified eIF4A1-CTD protein was concentrated to ∼12 mg/mL using a 3 kDa molecular weight cut off centrifugal concentrator, snap frozen in liquid nitrogen and stored at -80 °C.

### Generation of eIF4A1 proteins for SPR

cDNAs encoding recombinant N-terminal Avi- tagged human eIF4A1 full length (Avi-eIF4A1-FL) and eIF4A1 C-terminal domain (residues 237-406; Avi-eIF4A1-CTD) were synthesised by Genewiz (Suzhou, China) as *E. coli* codon optimized DNA sequences and cloned into the pBDDP-SPR3 vector containing an N- terminal double-His_8_ affinity tag followed by a TEV protease cleavage site.

For *in vivo* biotinylation, the proteins were expressed in *E. coli* CVB101 cells, previously lysogenized with the λ phage DE3 by using the λDE3 Lysogenization Kit (Merck). The cells were cultured in Terrific broth medium, supplemented with kanamycin and 50 μM biotin, and induced with 0.5 mM IPTG overnight at 18 °C. The cells were harvested by centrifugation and stored at -80 °C.

The cells were resuspended in Lysis buffer (50 mM Tris-HCl pH 8, 500 mM NaCl, 10% glycerol, 15 mM imidazole, 1.5 mM DTT, 1 mM PMSF, DNase I) and lysed by sonication. The lysates were clarified by centrifugation and applied to a 5 mL HisTrap FF column pre- equilibrated with IMAC buffer (50 mM Tris-HCl, pH 7.5, 500 mM NaCl, 5% glycerol) supplemented with 10 mM imidazole. After washing the column with IMAC buffer supplemented with 30 mM imidazole, the proteins were eluted with a linear gradient of 30- 500 mM imidazole in IMAC buffer, and subsequently incubated with TEV protease overnight at 4 °C for removal of the double-His_8_ tag. The TEV-treated samples were applied to a HiLoad 26/60 Superdex 200 pg size exclusion chromatography column pre-equilibrated with 20 mM Tris HCl pH 7.5, 500 mM NaCl, 5% glycerol, 1 mM DTT. The purified biotinylated Avi- eIF4A1-FL and Avi-eIF4A1-CTD proteins were concentrated to 1 and 21 mg/mL, respectively, using 10 kDa molecular weight cut off centrifugal concentrators, snap frozen in liquid nitrogen and stored at -80 °C.

### Surface plasmon resonance

For hippuristanol and epi-hippuristanol binding to eIF4A1, SPR experiments were conducted on a Biacore T200 (Cytiva) or Biacore 8K+ (Cytiva).

Purified, Avi-tagged eIF4A1 full length (Avi-eIF4A1-FL) and C-terminal domain, (Avi-eIF4A1- CTD) proteins were captured via the biotinylated avi-tag on Biacore Series S streptavidin sensor chips in running buffer (20 mM Tris pH7.5, 150 mM NaCl, 1 mM MgCl2, 1mM TCEP, 0.05% Tween20, 5% DMSO), to obtain capture levels of approximately 1000-1500 RU. Protein was captured on flow cells 2 and 4, with 1 and 3 used as reference surfaces (biotin blocked streptavidin) for the T200, and capture on FC2 with FC1 as reference on the 8K+.

Analysis and sample compartment temperatures were set to 25°C. Hippuristanol was tested in a single cycle kinetic assay format (300s contact time, 2400s dissociation), with a five-point titration, 3-fold concentration series from 10 µM,and a blank titration run both before and after.. Epi-hippuristanol was tested in a multicycle kinetic format (60s association, 60s dissociation) from 120 µM top concentration, with a 2-fold dilution series, 8-point titration. For all compound testing the flow rate was set to 30 µL/min. A solvent correction curve was run from 4% - 6% DMSO at the end of each run. Data A solvent correction curve was run from 4% - 6% DMSO at the end of the run. Data were analysed using Biacore Insight Evaluation software V5.0, (Cytiva). Solvent corrected sensorgrams were reference and blank subtracted with the blank run before compound titration, prior to fitting to a 1:1 binding model.

For eIF4G (MIF4G domain only) binding to eIF4A1, SPR experiments were performed as above, in a multi-cycle kinetic format, with the exception of capturing only 50- 100 RU eIF4A1 for protein interaction studies. 10x start up cycles with buffer were run to condition the protein prior to the eIF4G binding titration being run. Association was measured for 30s - 60s and dissociation for 60-120s. This experiment was repeated with either 10 µM hippuristanol or 10 µM eFT226 added to the running buffer, and the eIF4G-MIF protein samples for these runs were prepared in buffer containing compound. 10x start-up cycles were run prior to the eIF4G titration to saturate small molecule binding to eIF4A1.

### eIF4G-eIF4A1 dot blot interaction assay

A nitrocellulose membrane with pore size of 0.45 µM was inserted into a 96-well dot plot apparatus (BioRad). eIF4G-MC was then immobilised onto the membrane at 1.6 µg per well in 50 uL assay buffer (20 mM Hepes/KOH, pH 7.5, 100 mM KCl, 2 mM MgCl_2_). After the solution passed the membrane by gravitational flow, the membrane was blocked in 7% milk-TBST for 30 min. Serial dilutions of 0 – 3.6 ug eIF4A1 in 40 uL assay buffer supplemented with either 20 uM hippuristanol alone or 20 uM eFT266, 2 mM AMPNP and 50 nM (AG)_10_-RNA were applied to the wells and incubated until the solution passed the membrane. The membrane was then washed with TBST and developed using primary eIF4A1 (ab31217, Abcam UK) and secondary anti-rabbit (926-32213, LI-COR Biosciences).

### eIF4A1-CTD-hippuristanol low concentration complexing

400 µL of freeze-thawed, His- tag cleaved eIF4A1-CTD (12.1 mg mL^-1^, corresponding to 610 µM) was centrifuged for 5 mins at 13,000 g and 4C, then diluted to 25 µM in buffer B (20 mM Tris-HCl, pH 7.5, 300 mM NaCl, 5 % glycerol, 1 mM DTT, 5 % DMSO) to a final volume of 9.76 mL. 10 µL 100 mM hippuristanol (100 % DMSO), 190 µL buffer A (20 mM Tris-HCl, pH 7.5, 300 mM NaCl, 5 % glycerol, 1 mM DTT), 19.80 mL buffer B were mixed to give a final concentration of 50 µM hippuristanol in 5 % DMSO. eIF4A1-CTD (5 % DMSO) at 25 µM and hippuristanol (5 % DMSO) at 50 µM were mixed 1:1 (v/v) to give a final concentration of 12.5 µM eIF4A1 and 25 µM hippuristanol in 5 % DMSO, then incubated overnight at 4C. Subsequently, the sample was centrifuged for 5 mins at 4,000 g and 4C, and then re-concentrated to 11.3 mg mL^-1^ using 3 kDa MWCO spin concentrators pre-equilibrated with buffer B.

### Crystallisation, data collection and refinement

For apo-eIF4A1-CTD. eIF4A1-CTD at 12.1 mg mL^-1^ was crystallised using a fine screen based on preliminary crystals observed in SG1 condition G1 (16-30% (w/v) PEG8000, 0.2M sodium acetate trihydrate, 0.1M MES, pH 5.5-6.5) at 4C. A single crystal with needle-like morphology was flash-frozen in liquid nitrogen using 20% ethylene glycol as the cryoprotectant, and data were collected at Diamond Light Source (Harwell, UK) beamline I04, at 100 K and processed to 2.73 Å resolution using Xia2 (Winter, 2010). The structure was solved by molecular replacement using PHASER^37^ and the C-terminal domain of human eIF4A1 (residues 239-401; PDB 2ZU6)^38^ as the search model. The molecular replacement solution comprises 3 molecules in asymmetric unit (V_M_= 2.33 Å^3^ Da^-1^, 47.2% solvent content). The model was rebuilt using COOT^39^ and refined using REFMAC5^40^. The refinement statistics are shown in Supplementary Table 2.

For eIF4A1-CTD-hippuristanol, eIF4A1-CTD-hippuristanol (low concentration complexing) complex at 11.3 mg mL^-1^ was crystallised from PACT premier condition A7 (20 % (w/v) PEG 6000, 0.2 M sodium chloride, 0.1 M sodium acetate, pH 5.0) at 4C. A single crystal with needle-like morphology was flash-frozen in liquid nitrogen using 20% ethylene glycol as the cryoprotectant, and data were collected at Diamond Light Source (Harwell, UK) beamline I03, at 100 K and processed to 1.70 Å resolution using Xia2^41^. The structure was solved by molecular replacement using PHASER^37^ and the refined structure of human eIF4A1-CTD (PDB 9I9F) as the search model. The molecular replacement solution comprises 1 molecule in asymmetric unit (V_M_= 1.83 Å^3^ Da^-1^, 32.9% solvent content). The model was rebuilt using COOT^39^ and refined using REFMAC5^40^. The refinement statistics are shown in Supplementary Table 2.

The coordinates and structure factors for eIF4A1-CTD and eIF4A1-CTD-hippuristanol have been deposited at the PDB with accession codes 9I9Fand 9I9G respectively. Figures were prepared using PyMOL Version 2.5.8 (Schrödinger, LLC).

### Molecular dynamics

Each system was prepared using the AMBER ff14SB force field for the protein and the GAFF force field for the ligand. Ligand charges were assigned using Antechamber software at the AM1-BCC level of theory. The systems were solvated using the TIP3P water model, with a truncated octahedron box extending 10 Å beyond the solute. To neutralize the system, Na⁺ or Cl⁻ ions were added as appropriate.

Energy minimization was performed in two stages: first, with a restraint of 20 kcal·mol⁻¹·Å⁻² on the solute atoms using the conjugate gradient method, followed by a second minimization step without restraints. The system was then gradually heated to 300 K over 500 ps under constant volume conditions. This was followed by an equilibration phase at 1 atm constant pressure for 2 ns.

Production simulations were conducted for 200 ns, with snapshots saved every 100 ps, yielding a total of 1000 frames. Periodic boundary conditions were employed throughout the simulation, and the particle mesh Ewald (PME) method was used for long-range electrostatic interactions. Temperature control was maintained using a Langevin thermostat with a collision frequency of 5 ps⁻¹, while pressure was controlled using a Berendsen barostat with a relaxation time of 2 ps. The SHAKE algorithm was applied to constrain all hydrogen-involving bonds, allowing for a 2 fs timestep. A non-bonded interaction cutoff of 8 Å was applied. MD simulations were carried out using OpenMM (version 7.6)^42^.

### Sample preparation for Ribo-seq and Total RNA-seq

For one repeat experiment, MCF7 cells were seeded at 3x 5*10^6^ cells in 15 cm dishes per condition and per repeat experiment in DMEM (Gibco) supplemented with 10 % FBS (Gibco) and 2 mM final concentration of L-glutamine (Gibco). After 48 h, medium was replaced with fresh medium containing 150 nM hippuristanol or 20 nM eFT226. After 1 h of drug treatment, cells were harvested by scraping, washed in PBS, collected by centrifugation, snap frozen in liquid N_2_ and stored at - 80°C. A total of six repeat experiments were collected with cells seeded on different days.

For each biological repeat with organoids, a full 6-well plate with six plugs of organoids per well was used. For each repeat a different organoid line was used: lines 240614, 240615, RGH60.1g for eFT226, RVJ22.5c, RVJ22.5h, RVJ22.6d for hippuristanol experiments. Each cell pellet was lysed in 500 µl cold lysis buffer (15 mM Tris-HCl (pH 7.5), 15 mM MgCl_2_, 300 mM NaCl, 1% Triton X100, 0.05% Tween20, 2% n-Dodecyl-β Maltoside (Thermo Fisher 89903), 0.5 mM DDT, 100 µg/ml cycloheximide, 1X cOmplete EDTA-free Protease Inhibitor Cocktail (Roche 11836170001), 200 U/ml RiboLock RNase Inhibitor (Thermo Fisher Scientific, EO0381)), and cleared by centrifugation. For Total RNA-seq, 25 µl aliquots were taken and RNA extracted with TRIzol. For Ribo-seq, lysate were RNase-digested with 10 µl Ambion RNase I cloned, 100 U/µL (Thermo Fisher Scientific, AM2295) at 22°C for 15 min. Samples were loaded onto a 10-50% sucrose gradient (15 mM Tris-HCl (pH 7.5), 15 mM MgCl_2_, 300 mM NaCl and 100 µg/ml cycloheximide, BioComp gradient station) and were centrifuged at 38,000 rpm for 2 h at 4°C. Samples were then fractionated using a Biocomp gradient station and Gilson FC 203B fraction collector. 80S fractions were pooled, RNA extracted with acid phenol chloroform and separated on a 15% TBE-Urea gel (Thermo Fisher Scientific, EC68852BOX) for size selection of RPF fragments 28-34nt.

For RPF library preparation RNA was extracted and treated with T4 PNK for 5′ phosphorylation and 3′ dephosphorylation of the fragments. 10 ng of non rRNA-depleted RNA was input into the Nextflex small RNA v3 kit (Revvity, NOVA-5132-06), using the alternative step F bead clean up, 11 PCR cycles and gel extraction option.

For Total RNA-seq, taken RNA samples were checked for integrity and rRNA was depleted from 1 µg RNA with the RiboCop rRNA Depletion Kit HMR V2 (Lexogen 144.96).

Sequencing libraries were prepared using the Corall Total RNA-Seq Library Prep Kit (Lexogen 095.96) with 13 PCR cycles.

Final Total RNA-seq and Ribo-seq libraries were sequenced single-end on an Illumina NextSeq500 using a High Output v2/v2.5 kit (75 cycles).

### RNA-immunocoprecipitation of eIF4G (RIP seq) from MCF7 cells

About 3*10^7^ MCF7 cells per IP were harvested and lysed in lysis buffer (20 mM Tris pH 7.5, 150 mM NaCl, 40 mM KCl, 3 mM MgCl2, 0.5% Triton-X100, 1× protease inhibitors (Roche), 1 mM DTT, 1 mM NaF, 1 mM Na-β-glycerophosphate, 40 U/ml RiboLock (Thermo), 1% glycerol ). Lysates were cleared by centrifugation at 5000 rpm for 10 min at 4 °C. Aliquots of 60 uL were taken as input sample for total mRNA sequencing. Lysates were split equally and incubated with 3 ug anti eIF4G (Thermo Fisher Life Technologies, MA5-14971) or rabbit-IgG (Cell Signalling Technologies, #2729) antibodies for 60 min rotating at room temperature. Per IP, 40 uL Dynabeads™ M-280 Sheep Anti-Rabbit IgG were washed thrice with PBS, resuspended in 20 uL PBS and added to the lysate-antibody mix, allowing incubation for 30 min rotating at room temperature. After incubation beads were rinsed thrice with lysis buffer. At the final washing step, a quarter of resuspended beads was taken for downstream western blotting. The remaining three quarters were taken forward to RNA extraction. Beads for protein sample were resuspended in 1x SDS loading buffer, beads for RNA extraction were resuspended in 1 mL TRIzol. RNA was extracted and RNA libraries generated as described for the total RNA samples in the Ribo-seq section. Four repeat experiments for the RIP-Seq were performed with a different batch of cells grown on different days for each repeat. eIF4A1-inhibitors were present in all buffers essentially performing the IPs in the presence of the compounds.

### TMT-pSILAC

Experiments were performed as described previously^1^. Cells were treated with 20 nM eFT226. Data shown for treatment of MCF7 with hippuristanol are taken from our previous work^1^.

**NaP-TRAP and eIF4G-RIP-seq with NaP-TRAP library.**

All oligonucleotides and primers used in NaP-TRAP experiments are listed in **Supplementary Table 3**.

### Reporter construction

NaP-TRAP reporter libraries were constructed through PCR assembly using three different PCR reactions as mentioned in Strayer et al.^23^. For PCR1, the 3xFLAG-GFP fragment was amplified from the 3xFLAG-GFP plasmid (P63_F, P64_R). For PCR2, a custom single stranded DNA oligo pool consisting of 11,088 oligos (GenScript) was amplified using SP6 promoter and GFP-aug reverse (P61_F, and P62_R) primers^23^. Third, PCR three was divided into 2 steps. Step one connects the products of PCR1 and PCR2 by adding PCR1 and PCR2 products in equal molarity for 10 cycles. Step two amplifies the connected PCR products using SP6 forward and SV40 reverse primer with 60Ts to hard encode the poly(A) tail (P65_F and P113_R). All PCR reactions used the 2X KAPA HiFi HotStart ReadyMix (Roche #7958935001). All amplicons were gel purified (Zymo #D4001) and quantified using Nanodrop (Thermo Scientific #13-400-519).

### In vitro mRNA synthesis

To synthesize the NaP-TRAP reporter library, gel purified PCR3 product was used for in vitro transcription using a mMESSAGE mMACHINE™ SP6 transcription kit (mMESSAGE mMACHINE™ SP6 Transcription Kit #AM1340) following the manufacturer’s protocol. To remove the template DNA 1 uL of Turbo DNase (provided in the kit) was added in reaction mix and the reaction mix was incubated at 37C for 15 minutes. In vitro transcribed mRNAs were purified using a Zymo RNA clean and Concentrators (Zymo #R1013). The mRNA quality was analyzed by running it on 2% agarose gel and mRNA library was aliquoted and kept frozen at -80C.

### Library transfection and drugs treatment

The night before transfection, each well of a six well plates were seeded with 1.2 million MCF7 cells per well. The next morning, each well was transfected with 2.5 ug of library mRNA by diluting 2.5 ug of mRNAs in 125 uL of OptiMEM (Invitrogen™ #31985062) media and 2.5 uL of Lipofectamine messenger max (Thermo #LMRNA003) into 125 uL of OptiMEM (Invitrogen™ #31985062). Tubes were incubated at RT for 5 minutes. mRNA mix was added to lipofectamine mix and incubated for 5 minutes at RT. The mix was then added to the cells in a drop wise manner. Cells were incubated at 37C for one hour and washed with 1x PBS for three times to remove transfected mRNA. Cells were fed with fresh DMEM (Invitrogen™ #31985062) complete media, supplemented with 10% FBS (GeminiBio #100-106) and 2 mN L-Glutamine (Thermo #25030081) and incubated for an additional four hours. Post five hours transfection, the media was replaced with DMEM complete media supplemented with 20 nM eft226 or 150 nM hippuristanol or 0.1% DMSO (AmericanBio # AB03091-00100) and incubated for an additional 1 hour.

### Nascent-Peptide mediated Translating Ribosome Affinity Purification (NaP-TRAP)

For NaP-TRAP experiment, one well of a six well plate was considered as one replicate and total of four replicate were used per condition. NaP-TRAP was performed as detailed in Strayer et al. ^23^. Briefly, post six hours transfection, cells were treated with cycloheximide at a final concentration of 100 ug/mL and incubated at 37C for 10 minutes. Next, cells were kept on ice for 5 minutes and then washed with ice cold 1x PBS (supplemented with same concentration of respective drug or DMSO) for total of three times. Washed cells were lysed using 500 uL of low salt lysis buffer (15 mM Tris pH 7.5, 100 mM NaCl, 5 mM MgCl2, 1% Triton X100, 2 mM DTT, 1x Protease inhibitor, 100 ug/mL Cycloheximide, RNaseOUT 40U) and triturated using a 24Gz syringe at least 10 times. Lysate was transferred in a 1.5 mL low binding tubes and spun at 16,000g at 4C for 5 minutes, supernatants were transferred in new 1.5 mL low binding tubes. Next, 500 uL of high salt buffer (15 mM Tris pH 7.5, 1 M NaCl, 5 mM MgCl2, 1% Triton X100, 2 mM DTT, 1x Protease inhibitor, 100 ug/mL Cycloheximide, RNaseOUT 40U) was added to make up the volume to 1 mL and incubated with 6 units of DNaseI (NEB #M0303L) for 10 minutes on ice. During incubation the anti-FLAG magnetic beads (20 uL per sample) were washed with 800 uL of bead wash buffer (15 mM Tris pH 7.5, 500 mM NaCl, 5 mM MgCl2, 1% Triton X100, 2 mM DTT) with 15 seconds vortex and 30 seconds on magnetic stand (DynaMag™-2 Magnet #12321D) for a total of three times. Lysates were transferred in bead carrying, 1.5 mL low binding tubes and incubated for 2 hours at 4C with rotation. Post incubation, the magnetic beads were resuspended by gentle vortexing and 5% of input (50 uL) was collected in a new low biding 1.5 mL tube for each sample. The remaining beads were then washed with 800 uL of ice-cold wash buffer (15 mM Tris pH 7.5, 500 mM NaCl, 5 mM MgCl2, 1% Triton X100, 2 mM DTT, 1x Protease inhibitor, 100 ug/mL Cycloheximide, RNaseOUT 40U) with 15 seconds vortex and 30 seconds on magnetic stand for a total of three times. The mRNA-beads complex was finally resuspended in 1 mL of TRIzol (Thermo #15596018) supplemented with the 2 uL mix of different concentration of five different spike-ins.

For details of spike-ins design and construction please refer to Strayer et al.^23^. The spike-ins plasmids were a generous gift from the Giraldez lab. To generate spike-in mRNAs, plasmids were amplified with primers P65_F and P113_R (Table 1.1) using 2X KAPA HiFi HotStart ReadyMix (Roche #7958935001). The amplified product was gel purified (Zymo #D4001) and used for in vitro transcription with a mMESSAGE mMACHINE SP6 transcription kit (Invitrogen #AM1340). The mRNA was purified using a Zymo RNA clean and Concentrators (Zymo #R1013). The mRNA quality was analyzed by running it on a 2% agarose gel and five different spike-ins mRNA were pooled together in different concentration, 1, 5, 25, 50, and 125 fg (per 2 uL) for the 5’ UTR library. The pool of spike-ins was aliquoted and kept frozen at -80C.

**Table 1.** Kinetic parameters of compound binding to eIF4A1.

| parameter | eIF4A1-CTD |  | eIF4A1-FL |
| --- | --- | --- | --- |
|  | hippuristanol | epi-hippuristanol | hippuristanol |
| $K_D$ (nM) | $60.5 \pm 23.1$ | 27950 | $62.6 \pm 25$ |
| $k_{on}$ ($M^{-1} s^{-1}$) | $608 \pm 46.1$ | 5325 | $1000 \pm 138$ |
| $k_{off}$ ($s^{-1}$) | $3.97 \cdot 10^{-5} \pm 1.55 \cdot 10^{-5}$ | 0.149 | $6.55 \cdot 10^{-5} \pm 2.22 \cdot 10^{-5}$ |
| $\tau$ (s) | $29600 \pm 11900$ | n.d. | $16800 \pm 4680$ |
| note | note | n = 6, mean ± sd | n = 2, mean |

**Table 2.** Kinetic parameters eIF4G-MIF binding to eIF4A1.

| Compound | eIF4A1-FL |  |  | eIF4A1-CTD |  |
| --- | --- | --- | --- | --- | --- |
|  | DMSO | hipp | eFT226 | DMSO | hipp |
| $K_D$ (nM) | $21.3 \pm 7.6$ | $4.4 \pm 2.1$ | $29.6 \pm 21.8$ | > 10000 | $155 \pm 70$ |
| $k_{on}$ ($10^7 M^{-1} s^{-1}$) | $1.98 \pm 0.75$ | $2.5 \pm 0.8$ | $1.6 \pm 1.3$ | | |
| $k_{off}$ ($s^{-1}$) | $0.37 \pm 0.08$ | $0.11 \pm 0.02$ | $0.26 \pm 0.07$ | | |
| Repeats | n = 3 | n = 3 | n = 2 |  |  |

### eIF4G RNA immunoprecipitation sequencing (RIP-seq) on NaP-TRAP library

For RIP- seq experiment, two wells of a six well plate were considered as one replicate and such three replicates were used. The efficiency of the pulldown was measured using Western Blot. Post library transfection and drug treatment, cells were kept on ice for 5 minutes and each well was washed twice with 1x PBS supplemented with DMSO. Cells from each well were scrapped in 450 uL RIP-seq lysis buffer (20 mM Tris pH 7.5, 40 mM KCl, 150 mM NaCl, 3 mM MgCl2, 1 mM Na-β-glycerophosphate, 1 mM Sodium Flouride, 1 mM DTT, 1% Glycerol, 1% Trito X-100, 1x Protease inhibitors, 40U RNaseOUT (supplemented with 20 nM Zotatifin (eFT226) or 150 nM hippuristanol or DMSO) . Lysates from two wells were pooled to make one replicate. Lysates were transferred into fresh 1.5 mL low biding tubes. Cells were then triturated using a 24Gz syringe and left on ice for 15 minutes followed by centrifugation at 5200xg for 10 minutes at 4C. Supernatants were collected in a fresh low binding 1.5 mL tubes and 60 uL of lysates were collected as RNA inputs and 40 uL of lysates were collected as inputs for Western Blot. The remaining 800 uL supernatants were further divided into two tubes, 400 uL each, and incubated with either anti-eIF4G antibody (3ug) (Thermo Fisher Life Technologies #MA5-14971) or anti-Rabbit IgG (3ug) (Cell Signalling Technologies #2729) for 2 hours at 4C with rotation. While samples were incubating, for each IP reaction, 15 uL of anti-Rabbit Dynabeads™ M-280 Sheep Anti-Rabbit IgG (Thermo #11203D) beads were washed three times by adding 400 uL of 1x PBS supplemented with the same concentration of DMSO while beads are on magnetic stand (DynaMag™-2 Magnet #12321D) and finally diluted in 20 uL of 1XPBS. Post incubation, the beads were added to antibody-sample mix and incubated for 45 minutes at 4C and 30 minutes at room temperature with rotation. The beads were separated using magnetic stand (DynaMag™-2 Magnet #12321D) and 40 uL of flowthrough from each sample was collected as a control for Western Blot. The beads were washed with 400 uL of lysis buffer by gently vortexing the tubes for 5 seconds and then 1 minute on magnetic stand for a total of three times. During the final wash, 100 uL of beads solution was transferred after vortexing in a fresh 1.5 ml tube for Western Blot. Both tubes per sample, for RNA pulldown sequencing and Western Blot samples, were kept on magnetic stand and, upon bead separation, the supernatants were discarded, and 1 mL of TRIzol (Thermo #15596018) was added to the RNA pulldown beads and input sequencing samples prior to being stored at -80C. The pull-down beads for immunoblotting were resuspended in 14 uL of 1X SDS gel sample loading buffer while the input samples for immunoblotting were prepared by adding 13.5 uL of 4X gel loading buffer (BIO-RAD #1610747).

### RNA Isolation

For both NaP-TRAP and RIP-seq samples, TRIzol samples were thawed on ice and vortexed mixed. A 200 uL of chloroform (Sigma #CX1060-1) was added to each sample, vortexed for 15 seconds, and incubated on ice for 2 minutes. Samples were spun at 12,000g for 15 min at 4C. Around 480 uL of aqueous phase was transferred in a fresh 1.5 mL low binding tube having 2 uL of GlycoBlue (Thermo #AM9516) and briefly vortex mixed. Next, 500 uL of isopropanol (AmericanBio # AB07015-01000) was added to each sample, briefly vortexed and incubated on ice for 10 minutes. Post incubation, tubes were spun at max speed for 30 minutes at 4C. The supernatants were removed and pellets were washed with 1 ml of freshly prepared 75% ethanol by spinning the tube at 7,500xg for 5 minutes and then supernatants were removed. Similarly, pellets were washed again with 300 uL of 75% ethanol. Finally, tubes were spun again at high speed for 1 minute to remove excess of ethanol and RNA pellet was dissolved in 11 ul of nuclease free water (Sigma #W4502-1L).

### Reverse Transcription

cDNAs were synthesized using Superscript III (Thermo # Thermo # 18080044) following the manufacturer’s protocol and employing the same primer, Unique Molecular Identifier (UMI) and barcoding strategy as Stayer et al.^23^. Briefly, 1 uL of 10 mM dNTPs with 1 uL of reverse transcription primers (4 uM total for a mixed of staggered primers a/b, see Table 1.1) targeting the N-terminal of the 3xFLAG-GFP was added to each pulldown and input RNA samples. These primers contained a 6-nucleotide sample barcode and a 10 nucleotide UMI to allow for demultiplexing and removal of PCR duplicates, as well as a 3’ I7 Illumina adaptor sequence. To increase library complexity and facilitate sequencing, each set of primers barcode contained a pair of primer staggered by 1 nucleotide (e.g., RT1a and RT1b, see Table 1.1) . The mix tube was briefly spun and incubated at 65C for 5 minutes. Immediately, after incubation the tubes were kept on ice for at least 1 minute. Later, 4 uL 5X RT buffer, 1uL 100 mM DTT, 1uL 40U/uL RNaseOUT, and 1uL 200U/uL of Superscript III RT enzyme was added in each tube and pipette mixed. The tubes were briefly spun to collect all the material at bottom and incubated in thermal cycler at 55C for 1 hour. The RT enzyme was inactivated by incubating the tubes at 70C for 15 minutes. To remove untemplated RNA and RNA from RNA-DNA hybrid, a 3 uL mix of RNase H (NEB # M0297S) and RNas I (NEB # M0243S) was added to each sample and incubated at 37C for 15 minutes. The enzymes were inactivated by incubating the samples at 70C for 20 minutes.

### cDNA purification

Barcoded cDNAs of all replicates were pooled together for each sample and purified using AmpureXP beads (Beckman Coulter #A63880) following the manufacturer’s protocol. Briefly, 1.8x volume of Ampure XP beads were added in each sample and mixed 30 times using 200 uL pipette. The samples-bead mix was incubated at room temperature for 15 minutes. Post incubation, tube were kept on magnetic stand for 2 minutes, upon beads separation, supernatants were removed and beads were washed twice with 700 uL of freshly prepared 75% ethanol. The samples were air dried for ∼5 minutes to remove residual ethanol. The beads were resuspended in 20 uL of nuclease free water by brief vortexing and then pipetting 20 times. The tubes were incubated at room temperature for 2 minutes and then kept on magnetic stand for 2 minutes. Upon beads separation, elutes were transferred in separate 1.5 mL low binding tubes.

### Library Amplification and Purification

To limit PCR duplicates, a qPCR was performed to calculate the least number of PCR cycle required to amplify the library. Briefly, 10 ul of 2X KAPA HiFi HotStart ReadyMix (Roche #7958935001) mix with 1 uL of 10 uM Illumina i501 index, 1 uL of Illumina i701 index, 1 uL of purified cDNA, 0.8 uL of 2.5X Sybrgreen (Sigma #S9430), and 6.2 uL nuclease free water was added in one well of a qPCR plate. The qPCR was performed for 35 cycles of 98C, 65C, 72C for 10, 15, 20 seconds, respectively. Final amplification cycle was calculated by identifying the cycle number at half of max relative fluorescence unit (RFU) minus four cycles with 19 times more input cDNA. The libraries were then amplified using 2X KAPA HiFi HotStart ReadyMix (Roche #7958935001 with a combination of i50x and i70x series custom Illumina primers. The library was amplified using 0.3 uM of Illumina i5 series (i501, i502, i505, i506) and, 0.3 uM of Illumina i7 (i701, i702, i705, i706) series primers with 1X KAPA polymerase mix and 19 uL of purified cDNAs for the calculated number of cycles at 98C, 65C, 72C for 10, 15, and 20 seconds, respectively. The amplified library was purified using 1.8x volume of AmpureXP beads (Beckman Coulter #A63880) as described above.

### Library quantification and sequencing

The amplified libraries were quantified using TapeStations system (TapeStation 4200, Agilent Technology). Briefly, 1 uL of amplified libraries were mixed with 3 ul of D1000 sample preparation buffer (Agilent #5067-5602) and vortexed mixed for 1 minute. Next, samples were loaded onto D1000 screen tapes (Agilent #5067-5582) and ran in a TapeStation system (TapeStation 4200, Agilent Technology).

Sample quantification reports were generated and exported using system software. The library quantification was further validated by Qubit DNA HS Kit (Thermo # Q33230) using the manufacturer’s protocol. All samples were pooled together to have equal molar of each sample. The pooled library was further quantified using TapeStation D1000 screen tape and Qubit DNA HS kit. Finally, a 4nM dilution was prepared and sequenced on a Illumina NovaSeq 6000 platform.

### Bioinformatics analyses

#### NaP-TRAP and eIF4G RIP-seq of NaP-TRAP

Sequencing data were stored using LabxDB^43^. Read processing and mapping were performed as in Stayer et al.^23^. Paired-end reads were trimmed and demultiplexed using ReadKnead (https://github.com/vejnar/ReadKnead). Barcodes identifying replicates and UMIs were extracted from read two. The zebrafish 5’ UTR library reads were mapped to a library specific index using Bowtie v2.55. PCR duplicates were eliminated using UMIs. UMIs were considered identical if they had a Hamming Distance less than 2. Reads for each experiment were normalized by dividing the read counts of each reporter by the sum of the total number of reads mapped to the spike-ins. In the absence of spike-ins (e.g., RIP-seq experiments), read counts were normalized based on the total number of mapped reads per replicate (reads per million, RPM). Translation values were calculated as a ratio of reads in the pulldown relative to reads in the input. Translation values were only included in the downstream analyses if the input and corresponding pulldown contained greater than or equal to 15 unique reads across all replicates. All scripts used to process and analyze NaP-TRAP data were from Strayer et al.^23^ (https://github.com/ecstrayer/nap-trap_paper).”

### Ribo-seq and Total RNA-seq data processing and analysis

Data processing was performed using the https://github.com/Bushell-lab/Ribo-seq GitHub package and custom R scripts. Briefly, Ribo-seq 3’-adapter sequence 5’TGGAATTCTCGGGTGCCAAGG3’ and RNA-seq adapter sequence AGATCGGAAGAG were trimmed with cutadapt v4.2. Reads were deduplicated with umitools v1.1.2. For Ribo-seq reads, 4+4 nts UMIs flanking the RPF, for RNA-seq 12 nt UMIs were used. RNA-seq reads were aligned to protein coding transcripts (GCRCh38.p14) using bowtie2 v2.5.4. Ribo-seq reads were aligned with bbmap v38.98 to most abundant transcripts identified through the matching RNA-seq. Ribo-seq counts were quantified with custom python scripts, RNA-seq counts were quantified using rsem v1.3.3. Ribosomal occupancy reflects 5’UTR or CDS bound 80S excluding first 20 and last 10 codons.

### Genomes

*Human sapiens* GCRCh38.p14, *Mus musculus* GRCm39, Gallus gallus GRCg7b, D*anio rerio* GRCz11, D*rosophila melanogaster* BDGP6.46, *Arabidopsis thaliana* TAIR10, *Xenopus laevis* XenBase v10.1, *Caenorhabditis elegans* WBcel235, *Saccharomyces cerevisiae* S288C-R9.

### Alignments of eIF4A1 structures

AlphaFold2-predictions of eIF4A structures were downloaded from https://alphafold.ebi.ac.uk/, aligned using pyMol2 and the root mean square deviations were calculated by pyMol2.

### Differential expression analysis

Global differential expression of genes in ribosomal occupancy (RPF) or mRNA abundance (total RNA) alone or combined as translational efficiency (TE) was statistically examined using R package DESeq2 with RPF and total RNA counts as input. For RPF and total RNA the DESeq2 design *replicate + condition* was used, while the design *replicate + condition + seqtype + condition:seqtype* was used for TE. Standard adjusted p-values (0.1) and log2FC > 0 or log2FC < 0 were used to define up- and down-regulated targets, respectively.

### Gene set enrichment analysis

The R package fgsea was used for gene set enrichment analyses (GSEA). The log2FC in translational efficiency were used as input with the human or mouse specific pathway reference lists. Output terms were filter for adjusted p-values < 0.1.

### Gradient boosting

Gradient boosting was performed using the R package *gbm* v2.2.2. An input table of mRNA features of human and mouse mRNAs was generated by calculation of parameters through the R package *seqinr*. Features that correlated highly with the changes in translational efficiency (|R| > 0.7) were removed from the analysis. For regression of experimental data the log2FC in translational efficiency following hippuristanol or eFT226- treatment were filtered for approximately equal group sizes of sensitive and insensitive genes based on their adjusted p-value of the DESeq2 analysis. Data for human and mouse were subsequently joined, retaining the mouse and human specific transcript information i.e., per gene symbol the most abundant human and mouse transcript were retained. The table was then split into train and test datasets (0.7/0.3) before gradient boosting. Resulting models were quality controlled via AUC/ROC analysis and their accuracy in predicting the experimental data tested via Pearson correlation against the test dataset using the *caret* and *ROCR* packages. Best model parameters for EFT226: shrinkage = 0.03, interaction.depth = 30, n.minobsinnode = 3, bag.fraction = 0.7, optimal_trees = 167; HIPP: shrinkage = 0.01, interaction.depth = 40, n.minobsinnode = 5, bag.fraction = 0.7, optimal_trees = 308.

### Differential motif enrichment analysis

The webserver of the MEME-suite STREME was utilised to examine differentially enriched 5’UTR motifs of eFT226-sensitive over hippuristanol-sensitive (or *vice versa*) or over insensitive mRNAs. The top three discovered motifs with E-values < 0.1 are shown.

### Relative Synonymous Codon Usage

The relative synonymous codon usage (RSCU) was calculated using the *seqinr* R package.

### RIP-seq analysis

Sequencing data were processed following standard pipeline for total RNA sequencing. To statistically test for specifically enriched and depleted mRNAs bound by eIF4G over the input, DESeq2 was utilised using the following design *replicate + condition + seqtype + condition:seqtype.* Seqtype: RIP or input, condition: DMSO, eFT226 or hippuristanol. Standard adjusted p-values (0.1) and log2FCs ( |x|>0) were used.

### uORF identification

Alternative open reading frames were identified and annotated using standard parameters within the *ribotricer* package with raw sequencing data as input and the GCRCh38.p14 genome assembly as reference. Annotations of output ORFs were fed into the Ribo-seq pipeline as stated under section Ribo-seq and Total RNA-seq data processing and analysis to generate count files for further analysis.

### Evolution analysis

The following reference genomes were used for the analysis: GRCg7b (*Gallus gallus*), GCRm39 (*Mus musculus*), GCRCh38.p14 (*Homo sapiens*), GRCz11 (*Danio rerio*), BDGP6.46 (*Drosophila melanogaster*), TAIR10 (*Arabidopsis thaliana*), XenBase v10.1 (*Xenopus leavis*), WBcel235 (*Caenorhabditis elegans*), S288C-R9 (*Saccharomyces cerevisiae*). Genomes were filtered for length outliers and positional conservation of 5’UTRs and coding regions across species was scored using the *Biostrings* R package. Human orthologs were annotated as given in the respective genome databases. Feature detection and analysis was performed as stated in above sections using the individual genome files of the different species. *S. cerevisae* and *D. melanogaster* eIF4G RIP-seq data were taken from GEO GSE97438^44^ and Array express E-MTAB-2464^45^.

### Statistical analysis

n is the number of independent repeat experiments (not technical repeats) of the described experiment and is given in the figure legends. In of sequencing data generated from mouse organoids, n is the number of biological repeats (organoids derived from different animals). Statistical tests used are stated in the figure legends. Where applicable p-values were corrected for multiple testing by calculating FDRs. Statistical significances are given as the absolute, adjusted p-values in the figures or figure legends.

### BioRender license information

In the format https://BioRender.com/figure-specific-code Figure 1: t54q303, Figure 2: x25k751, Figure 3: w42i304, Figure 4: a37t546, Figure 6: v37q628, Supplementary Figure 2: e64n071, Supplementary Figure 3: s16o256, Supplementary Figure 4: d99g336, Supplementary Figure 7: u21p974

### Data and Code Availability

Custom R and python scripts that have been used to generate data tables and figures from sequencing and proteomics data can be found at https://github.com/ecstrayer/nap-trap_paper and https://github.com/Bushell-lab/.

Result data tables regarding differential RNA abundance, ribosomal occupancy, protein abundance, translational efficiency, reporter translation and eIF4G-binding are deposited and accessible together with the respective raw data below. The Ribo-seq in MCF7 cells and mouse organoids derived from a colorectal cancer model (*VillinCre*^ER^ *Apc^-^*^/-^, *Kras*^G12D/+^) are deposited at GEO under accession numbers GSE284484 and GSE284480, respectively. The eIF4G RIP-seq from MCF7 cells is available at GEO with accession number GSE284477.

Sequencing data from NaP-TRAP reporter translation and NaP-TRAP eIF4G RIP-seq are deposited at GEO under accession numbers GSE289358 and GSE289359, respectively. The TMT-pulsed SILAC mass spectrometry data have been deposited to the ProteomeXchange Consortium via the PRIDE ^46^ partner repository with the dataset identifier PXD058531 The coordinates and structure factors for eIF4A1-CTD and eIF4A1-CTD-hippuristanol have been deposited at the PDB with accession codes 9I9F and 9I9G respectively.

## Acknowledgements

We would like to thank the Core Services and Advanced Technologies at the Cancer Research UK Scotland Institute (C596/A17196 and A31287), with particular thanks to the Beatson Molecular Technologies team. We thank Sergio Lilla for handling the deposition of the TMT-pSILAC data to the ProteomeXchange Consortium. We thank Meghan Thomson for supporting the crystallography trials, and Emma Carswell and Laurent J.M. Rigoreau for directing the crystallography experiments and discussion of the data presented within this manuscript. The manuscript was critically reviewed by Catherine Winchester (CRUK Scotland Institute).

## Author Contributions

T.S. conceived and designed the project, designed and performed ribosome-profiling and TMT-pSILAC, eIF4G-RIP-seq, experiments, computational data analysis including NaP-TRAP and evolution, created figures, wrote the manuscript (draft and edits), and supported acquisition of funding.T.S. produced and purified recombinant proteins and performed, analysed and interpreted *in vitro* ATPase, RNA-binding and unwinding as well as *in vitro* translation assays. A.P.T. performed the structural studies, analysis, created structure images and contributed to writing the manuscript. A.G. designed and performed NaP-TRAP and NaP-TRAP eIF4G RIP-seq experiments. J.A.W. optimised and performed organoid ribosome-profiling in colorectal cancer mouse models and provided R- and python-code to support the analysis of Ribosome-profiling data. K.P. performed, analysed, prepared data tables and plots of SPR experiments. J.M. prepared sequencing libraries for ribosome profiling from eFT226 treated organoids. R.N. performed and prepared data plots from molecular dynamics simulations. P.H. wrote R- and python-scripts to annotate uORFs identified with ribotricer. A.D. expressed and purified recombinant proteins and performed and analysed RNA-binding, ATPase and unwinding assays. K.H. performed mass spectrometry-related sample processing, mass spectrometry and raw data processing. A.M.V. processed and analyzed NaP-TRAP data. L.P. and C.P. produced protein for crystallography and SPR. G.K. generated and cultivated mouse organoids derived from mouse small intestines. S.Z and O.J.S. supervised experiments, contributed to discussion and provided resources. J.-D.B. designed, performed and processed and analysed data from NaP-TRAP and NaP-TRAP eIF4G RIP-seq experiments. M.B. conceived and designed the project, designed experiments, assisted with manuscript writing, edit the manuscript and acquired funding. All authors gave feedback on the manscript and approved the final version.

## Supplementary Figures

**Supplementary Figure 1.**
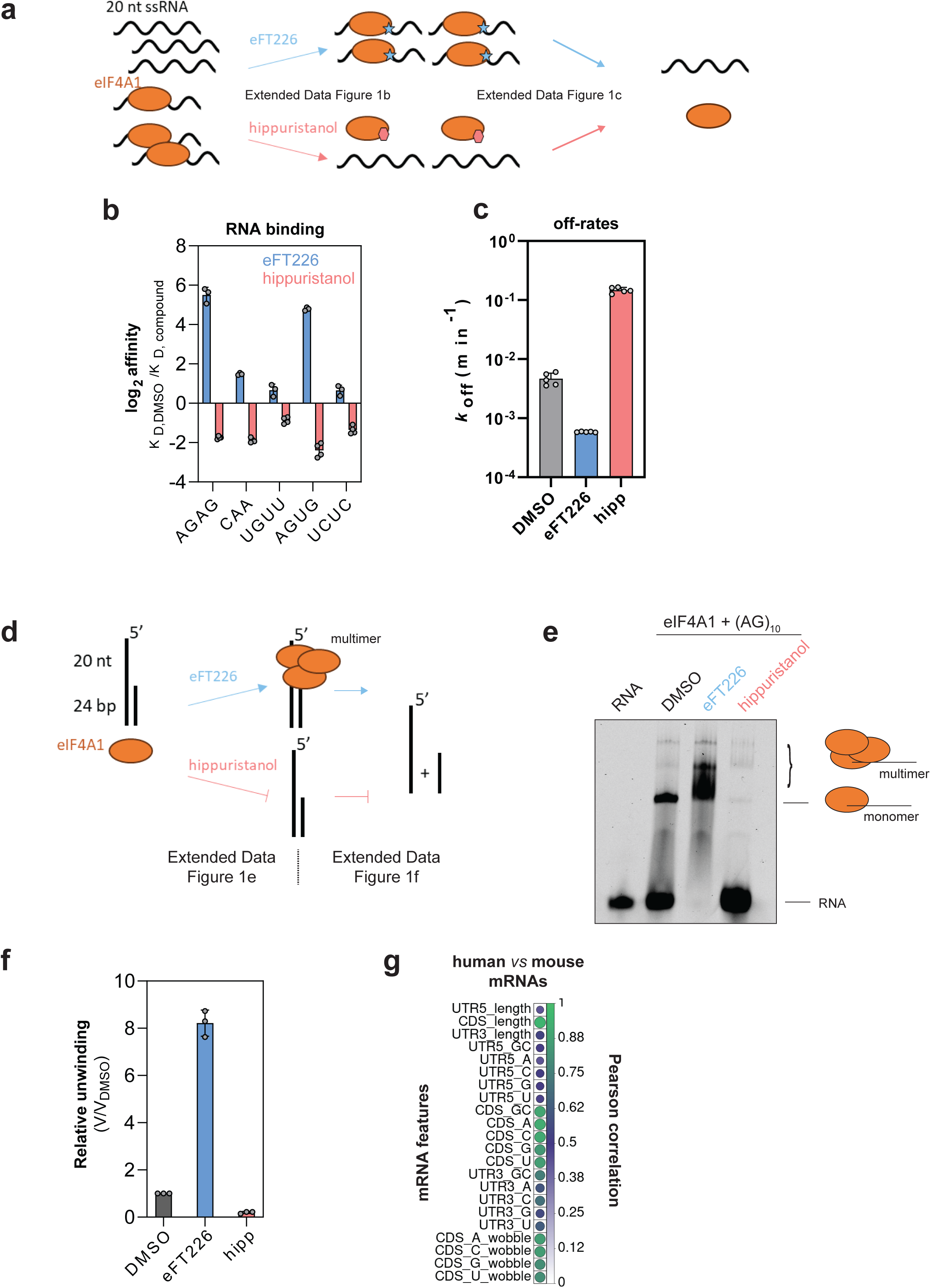
**a**, Schematic summary of the effects of eFT226- and hippuristanol-binding to eIF4A1 on RNA-binding by eIF4A1. **b**, log2-fold changes in RNA- binding (K_D_ – dissociation constant) of eIF4A1 to indicated 20 nt repeat ssRNAs in the presence of eFT226 or hippuristanol. Data are mean ± sd from four repeat experiments. **c**, Off-rates of RNA release by eIF4A1. Data are mean ± sd from five repeat experiments. **d**, Schematic summary of the effects of eFT226- and hippuristanol-binding to eIF4A1 on eIF4A1-multimerisation and RNA unwinding by eIF4A1. **e**, Electrophoretic mobility shift assay of eIF4A1 binding to AG-repeat ssRNA in the presence of AMPPNP and eFT226 or hippuristanol, respectively. **f**, Relative unwinding of an RNA substrate with 20 nt AG-repeat 5’ overhang and 24bp duplex region (see panel 1d) by eIF4A1 in the presence of eFT226 or hippuristanol. Data are mean ± sd from three repeat experiments. **g**, Pearson correlation coefficients between indicated mRNA sequence features of human and mouse transcripts.

**Supplementary Figure 2.**
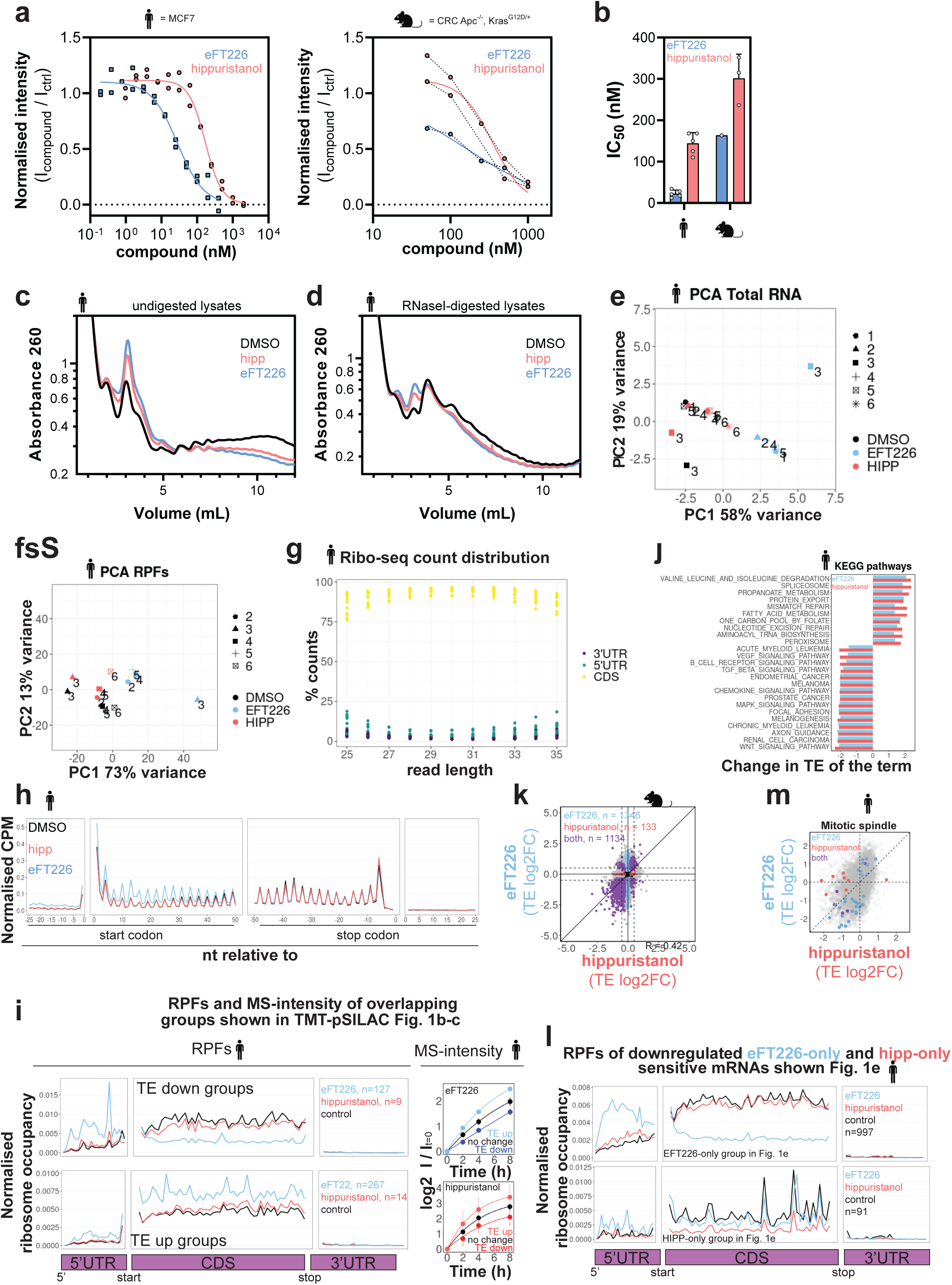
**a**, Representative dose-response curves of global translation of MCF7 cells and **b**, mouse organoids (CRC, Apc^-/-^/Kras^G12D/+^) following incubation with indicated compound concentrations for one hour measured by puromycin incorporation. For mouse eFT226 a single measurement is shown, for the other conditions in panels 2a-b a technical duplicate is shown. solid line – data fit, dashed line – connection between dots per replicate. **b**, Quantification of IC_50_ values of n repeat experiments of experiments shown representatively in a and b. Data are mean ± sd, n(MCF7) = 5; n(mouse, eFT226)=1, n(mouse, hippuristanol) = 3. **c**, UV traces of polysome gradients of undigested and **d**, RNase-treated MCF7 cell lysates following cell treatment with IC_50_-concentrations of eFT226 or hippuristanol for one hour. The mean of four repeats is shown. **e**, Principal component analysis of batch-corrected RNA-seq and **f**, Ribo-seq samples from MCF7 experiments. **g**, Distribution of read counts of ribosome protected fragments (RPF) from the Ribo-seq between 5’UTR, CDS and 3’UTR in transcripts across read lengths from MCF7 experiments for each replicate and condition (4 repeats x 3 conditions = 12 datapoints per region). **h**, metagene presentation of mean counts per million of RPF reads of all samples per condition from MCF7 experiments. **i,** Metagene analysis of the mean ribosome occupancy across the whole mRNA (left panels) and heavy-label TMT-pSILAC intensities (i.e., protein synthesis) normalised to t = 0 h of translationally down- and up-regulated genes that are identified in both Ribo-seq and TMT-pulsed SILAC. Grouping is the same as shown in pSILAC figure 1b- c. **j**, KEGG pathway enrichment analysis based on changes in TE of genes following hippuristanol or eFT226 treatment of MCF7 cells. **k,** Mean log2-fold changes in translational efficiencies (TE) per gene following eFT226 and hippuristanol treatment in mouse organoids. R – Pearson correlation coefficient. **l,** Metagene analysis of the mean ribosome occupancy across the whole mRNA of translationally downregulated genes that are only sensitive to either eFT226 or hippuristanol. Grouping is the same as shown in figure 1e for hippuristanol and eFT226. **m**, Mean log2-fold changes in TE per gene of the GO terms mitotic spindle and WNT signaling following hippuristanol or eFT226 treatment of MCF7 cells.

**Supplementary Figure 3.**
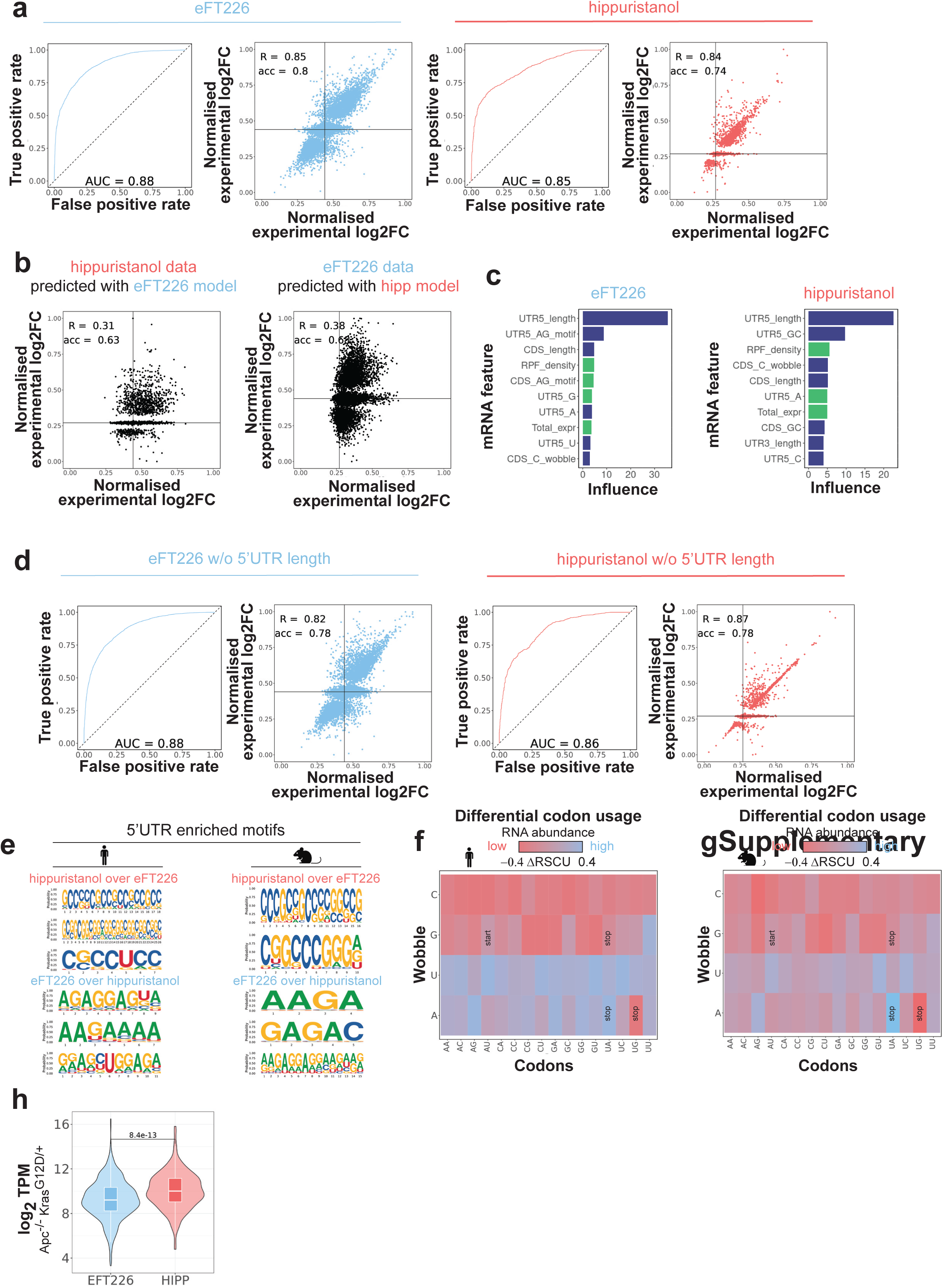
**a**, ROC curves of the gradient boosting models (including 5’UTR length) for eFT226 and hippuristanol data together with predicted log2-fold changes in translational efficiency plotted against the experimental data. R - Pearson correlation between predicted and experimental log2-fold changes in TE. Accuracy – Precision in predicting compound sensitivity tested again the input data. **b**, Prediction of hippuristanol experimental log2-fold changes in translational efficiency using the eFT226 gradient boosting model and *vice versa*. R - Pearson correlation between predicted and experimental log2-fold changes in TE. Accuracy – Precision in predicting compound sensitivity tested again the input data. **c**, Top ten mRNA features (including 5’UTR length) most influential in predicting mRNA sensitivity to eFT226 or **c**, hippuristanol as derived by gradient boosting. Input data combined from human and mouse Ribo-seq. **d**, ROC curves of the gradient boosting models excluding 5’UTR length for eFT226 and hippuristanol data together with predicted log2-fold changes in translational efficiency plotted against the experimental data. R - Pearson correlation between predicted and experimental log2-fold changes in TE. Accuracy – Precision in predicting compound sensitivity tested again the input data. **e**, Differentially enriched 5’UTR sequence motifs of hippuristanol- *vs* eFT226-*down* or vice versa for human and mouse target mRNAs individually. **f-g**, Differential synonymous codon usage of the 10% most and least abundant mRNAs in MCF7 (f) and mouse organoids (g). **h**, Transcript abundance (log2 TPM) from tissue RNA-seq data taken from Vande Voorde *et al*.^47^ P-value calculated from Wilcoxon-test. Box plots show median ± interquartile range.

**Supplementary Figure 4.**
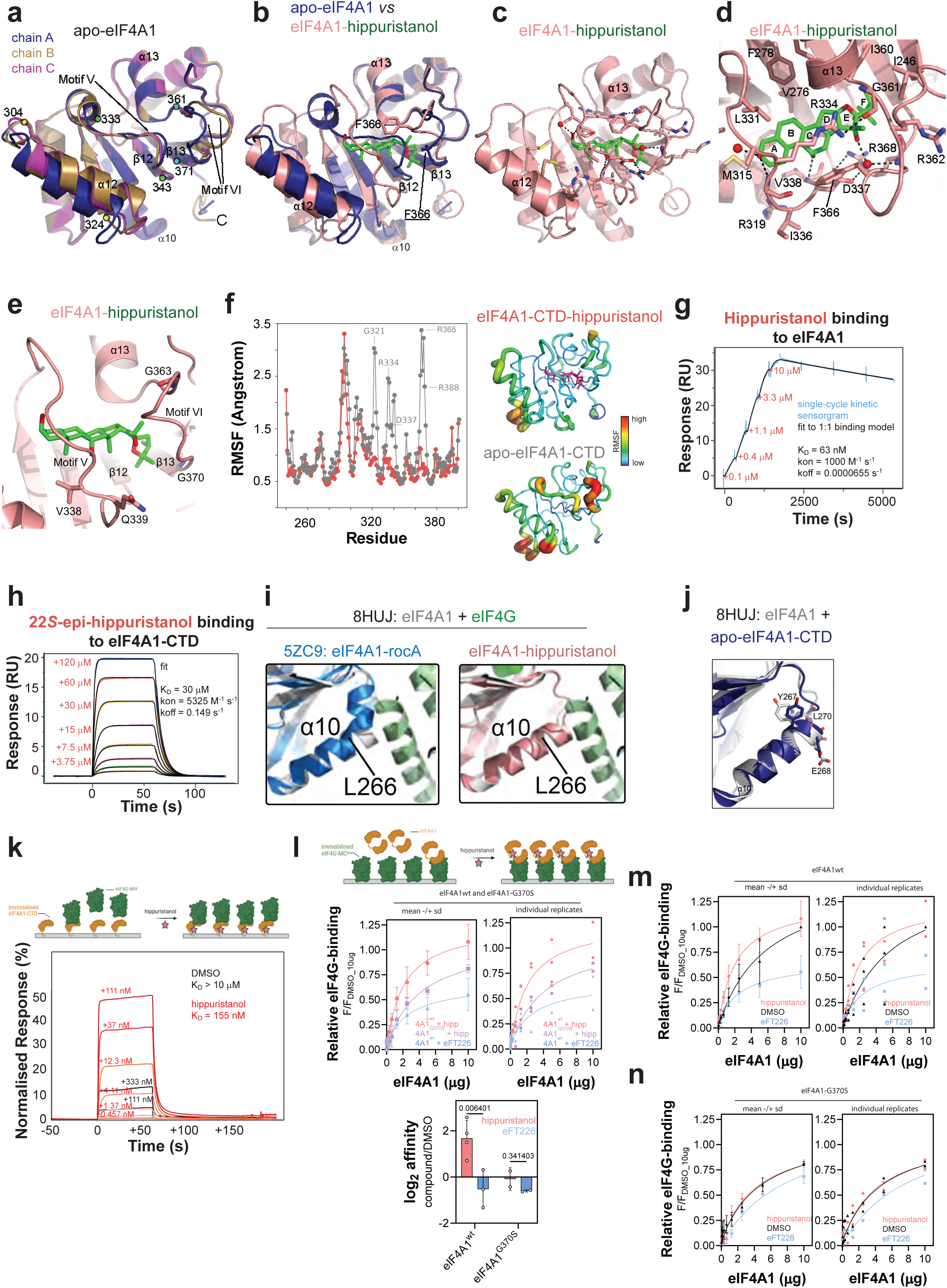
**a**, Overall structure of apo-eIF4A1-CTD (PDBid 9I9F). Superposition of the three chains in the asymmetric unit coloured dark blue (chain A), brown (chain B), and magenta (chain C), respectively. Structural regions displaying the largest differences in conformation between the three chains correspond to residues 304-324 (helix α12), 333-343 (motif V loop), and 361-371 (motif VI loop connecting α13 and β13; residues 364 to 367 inclusive are disordered in chain C), and are denoted by coloured dots. **b**, Superposition of eIF4A1-CTD in complex with hippuristanol (red) on apo-eIF4A1-CTD chain A (dark blue). Hippuristanol is shown with carbon atoms colored green. Hippuristanol displaces the motif V loop, with concomitant unwinding of residues D330 to L332, and the motif VI loop. Compared with apo-eIF4A1-CTD, motif VI in the eIF4A1-CTD-hippuristanol complex adopts a conformation in which F366 (labelled) is pointing inwards and flanking the bound hippuristanol molecule, whereas in apo-eIF4A1-CTD, the motif VI loop adopts a conformation where F366 is pointing outwards (chain B; F366 labelled and underlined) or is disordered (chains A and C). In addition, α12 is further displaced in the eIF4A1-CTD- hippuristanol complex. **c**, eIF4A1-CTD (red) in complex with hippuristanol (green) highlighting residues within 5 Å of hippuristanol shown as a stick representation. Water molecules are represented as red spheres. Hydrogen-bonding interactions to hippuristanol are represented as dashed black lines. **d**, rotation of the view in main figure 3b by 75° on the x-axis. Hippuristanol rings A to F are labelled. Residues with an atom within 5 Å of hippuristanol are labelled. Hydrogen-bonding interactions are represented as dashed black lines. Water molecules are represented as red spheres. **e**, eIF4A1-CTD bound to hippuristanol highlighting residues, mutations of which were previously shown to confer hippuristanol resistance^2,12^. **f**, Root mean squared fluctuation (RMSF) calculation from trajectories of 400 ns using classical MD simulations of hippuristanol bound to eIF4A1-CTD (red) or apo-eIF4A1 alone (grey). In the structures the respective RMSF is mapped on the protein domain structure. **g**, Representative single-cycle SPR-sensogramm of hippuristanol binding to immobilised, full-length avi-tagged eIF4A1. Concentrations given in the plot indicate time points of hippuristanol injections. Three repeat experiments have been performed. **h**, Representative, multi-cycle SPR-sensorgram of 22*S*-epi-hippuristanol binding to immobilised avi-tagged eIF4A1-CTD. Concentrations of 22*S*-epi-hippuristanol are given in the plot. Two repeat experiments have been performed. **i**, comparison of structures shown in Figure 3h with a cryo-EM-structure of the eIF4A1-eIF4G complex (green, PDBid 8huj) focusing on the conformation of α10. **j**, eIF4A1-CTD from apo-eIF4A1 (blue, chain A) superimposed on eIF4A1 from the eIF4A1- eIF4G complex (grey, PDBid 8HUJ) highlighting the positions of residues implicated in hydrogen bonding interactions in the eIF4A1-eIF4G complex. **k**, Representative, multi-cycle SPR-sensorgram (normalised to R_max_ = 100%) of indicated concentrations of eIF4G-MIF binding to immobilised, avi-tagged eIF4A1-CTD in the absence and presence of 10 µM hippuristanol. Three repeat experiments have been performed and are summarised in Table 2. **l**, Titration of eIF4A1^wt^ and hippuristanol-resistant eIF4A1^G370S^ to immobilised eIF4G-MC in the presence of hippuristanol or eFT226, and quantification of apparent log2-fold change of the binding constant. p-value calculated with paired t-test. Data are mean ± sd from n(eIF4A1^wt^+hipp) = 4, n(eIF4A1^wt^+eFT226) = 3, n(eIF4A1^G370S^) = 2 repeat experiments. For individual replicates, see next panels. **m-n,** Titration of eIF4A1^wt^ and hippuristanol-resistant eIF4A1^G370S^ to immobilised eIF4G-MC in the presence of hippuristanol or eFT226, and quantification of apparent log2-fold change of the binding constant. Data show individual data points or mean ± sd from n(eIF4A1^wt^+hipp) = 4, n(eIF4A1^wt^+eFT226) = 3, n(eIF4A1^G370S^) = 2 repeat experiments.

**Supplementary Figure 5.**
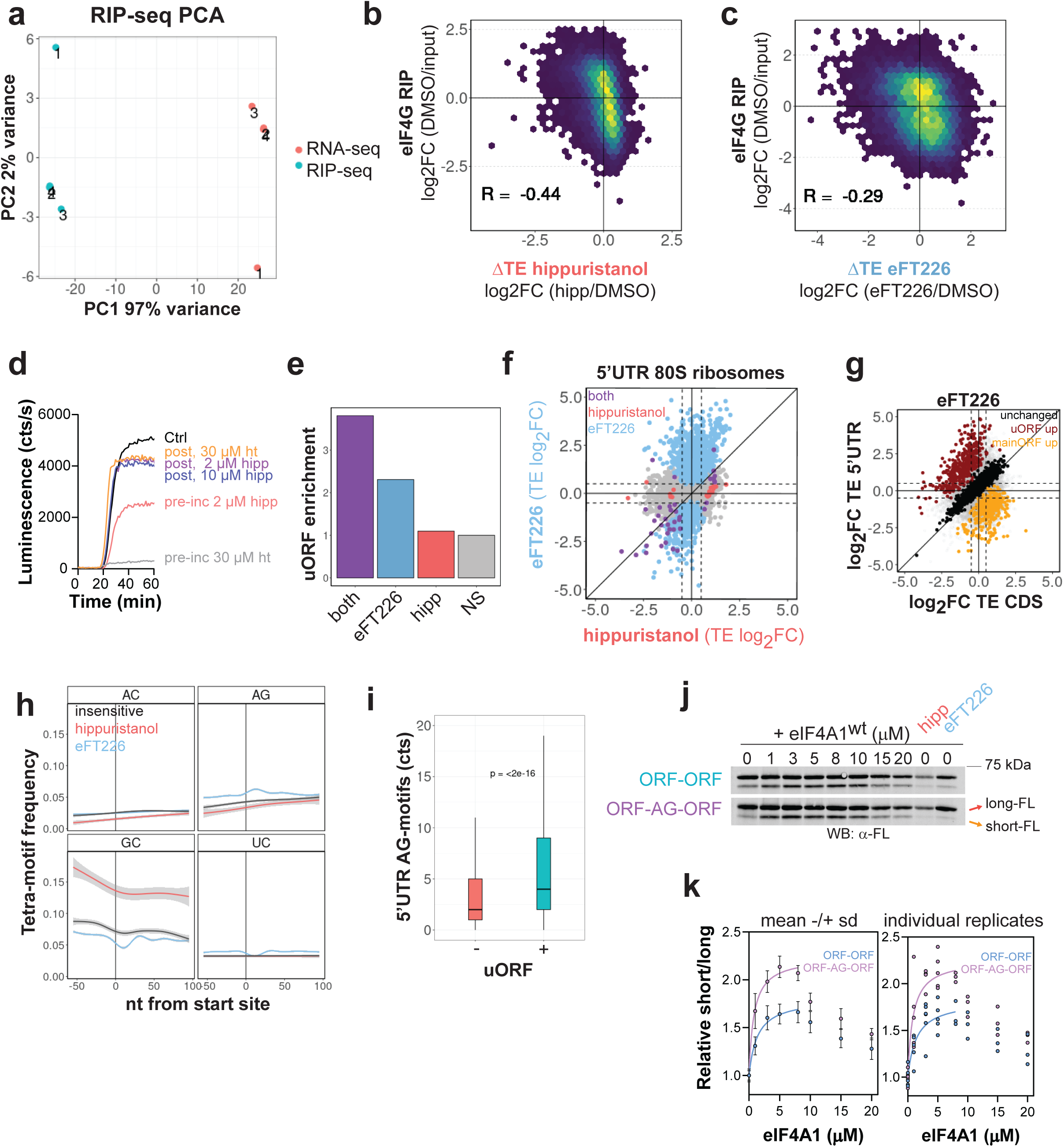
**a**, Principal component analysis of batch-corrected eIF4G RIP-seq samples from MCF7 cells. **b**, 2D density plot between mean enrichment in eIF4G- binding (eIF4G-RIP-seq) and mean log2-fold changes in translational efficiency following hippuristanol or **c**, eFT226 treatment (Ribo-seq). **d**, Real-time progress curves of firefly luciferase activity from reporter translation in rabbit reticulocyte lysate with indicated incubation scheme of the addition of hippuristanol (time points in legend) and reporter mRNA (t = 0 min). hipp – hippuristanol, ht - harringtonine **e**, Enrichment of mRNAs containing uORFs of eFT226- and hippuristanol-sensitive mRNAs relative to insensitive mRNAs. **f**, Mean log2-fold changes in translational efficiencies (TE) in the 5’UTR of genes following eFT226 and hippuristanol treatment. Data reflects changes in 80S ribosomal occupancy within the 5’UTRs of transcripts. **g**, Mean log2-fold changes in translational efficiencies (TE) in the 5’UTR and coding region of genes following eFT226 treatment; calculated from Ribo-seq data shown in Figure 1c. **h**, Frequency of indicated 4mer-motifs downstream of alternative start sites within the 5’UTR of eFT226-, hippuristanol-sensitive or compound- insensitive mRNAs. Data are mean ± 95% ci. **i**, Number of 5’UTR AG-motifs in mRNAs with (+) or without (-) uORFs within their 5’UTRs. Box plot shows median ± interquartile range. P-value calculated from Wilcoxon-test. **j**, Representative western blot of *in vitro* translation reactions of reporters shown in main figure 4j in RRL detecting firefly luciferase production. **k**, Quantification of data representatively show in panel 5**j**; data are mean ± sd, n (eIF4A1 < 8 µM) = 5, n (eIF4A1 ≥ 8 µM) = 3.

**Supplementary Figure 6.**
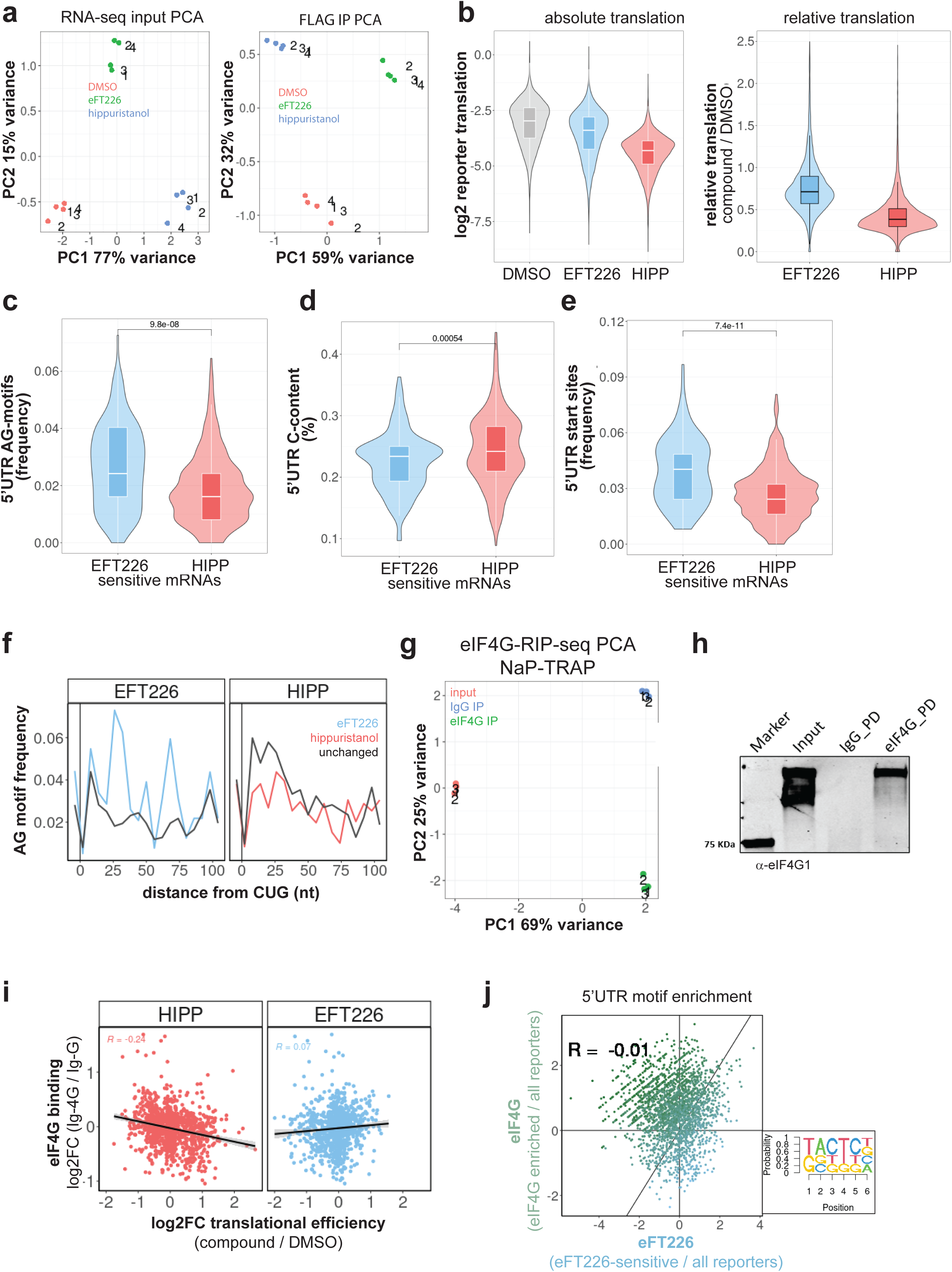
**a**, Principal component analysis of batch-corrected RNA-seq from input and FLAG-IP samples from the NaP-TRAP assay. **b**, Mean log2-translation values (IP/input, left) and translational values relative to DMSO (right) of spike-in corrected reporter mRNAs following DMSO or compound-treatment. Box plot shows median ± interquartile range. **c-e**, Difference in indicated 5’UTR features between hippuristanol and eFT226- sensitive reporters. P-values calculated from Wilcoxon-test. Box plot shows median ± interquartile range. **f**, Frequency of AG-motifs within the 5’UTR of reporter mRNAs downstream of CTG start sites of hippuristanol- and eFT226-inhibted reporters compared to insensitive mRNA reporters. **g**, Principal component analysis of batch-corrected RNA-seq from input and eIF4G-IP and IgG-IP samples from the eIF4G-RIP-seq of the NaP-TRAP reporter library. **h**, Representative western blot of eIF4G- and IgG-control pulldowns (PD) from NaP-TRAP assay used for RIP-seq. **i**, Mean log2-fold change in translational efficiency of reporters following treatment with eFT226 or hippuristanol plotted against their log2- enrichment in the eIF4G-RIP-seq under control DMSO conditions (eIF4G-IP / IgG-IP). R – Pearson correlation coefficient. Black line is a linear fit to the data ± 95% ci. **j**, Log2- enrichment (eFT226-sensitive or eIF4G-enriched reporters / all reporters) of 5-6mer sequence motifs within the 5’UTRs of eFT226-sensitive reporters and reporters enriched for eIF4G binding. R – Pearson correlation coefficient.

**Supplementary Figure 7.**
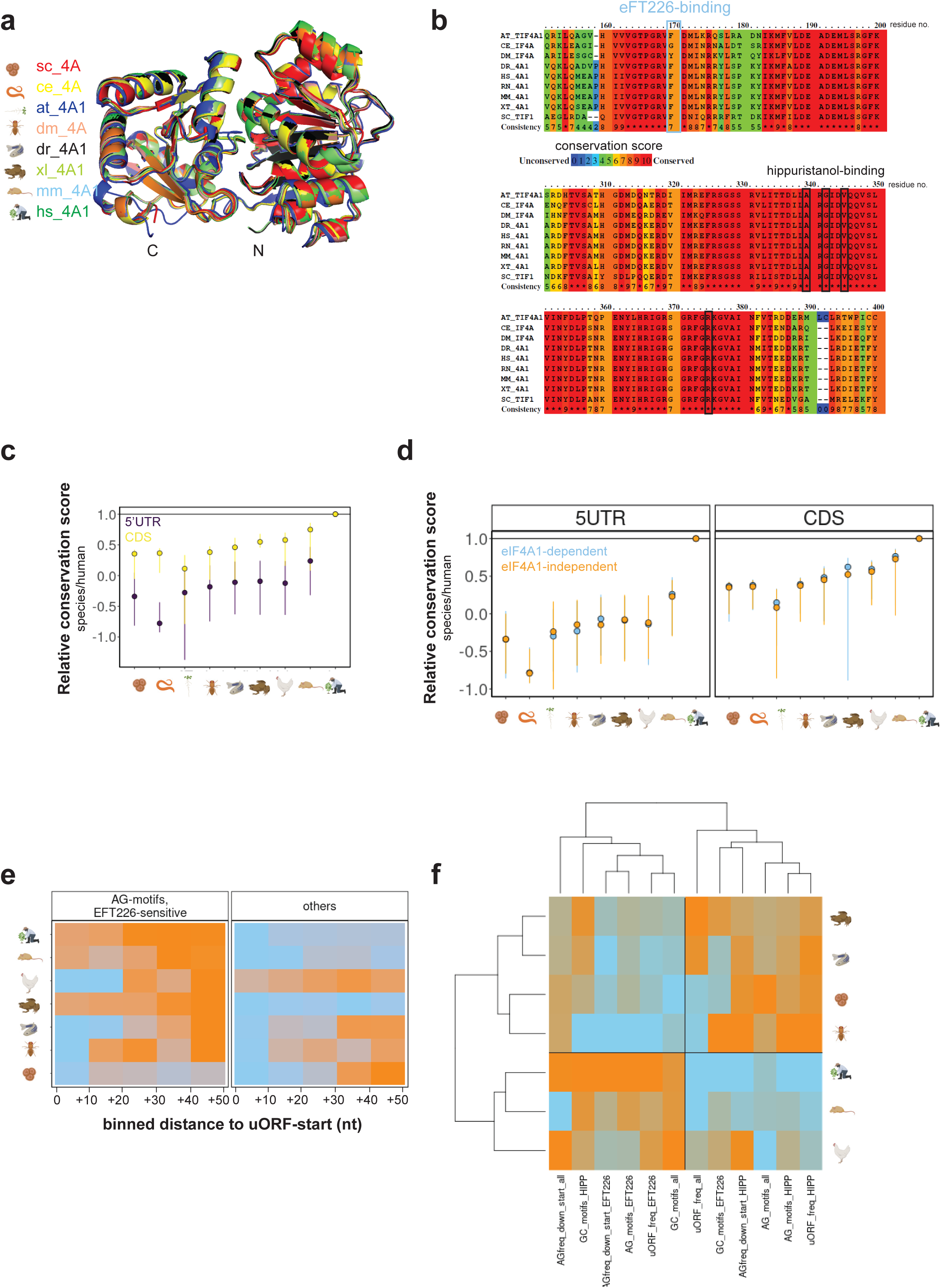
**a**, Superposition of indicated, Alphafold2-predicted structure of eIF4A proteins from different species. N- and C-termini are labelled “N” and “C”. **b**, Alignment of the primary sequence of eIF4A from indicated species. The key residue responsible for eFT226-binding, corresponding to F163 in human eIF4A1, is highlighted in blue. The key residues responsible for hippuristanol-binding, corresponding to A333, G335, V338 and R368 in human eIF4A1, are highlighted in black. **c**, Transcriptome-wide conservation scores of the 5’UTRs and CDSs of indicated species relative to human. Data are median ± interquartile range (iqr). **d**, Conservation scores of the 5’UTRs and CDSs of eIF4A1-dependent and -independent homologue mRNAs of indicated species relative to human. Data are median ± iqr. **e**, Frequency of AG-motifs in 10-nt bins downstream of alternative start sites in 5’UTRs of indicated species normalised within each species for eFT226-sensitive and all other transcripts. **f**, Non-supervised clustering of data shown in main Figure 6d.

## Supplementary Movie

Movie showing the trajectory of 400 ns using funnel MD simulation of the eIF4A1-CTD- hippuristanol complex.

## Supplementary Tables

**Supplementary Table 1.**
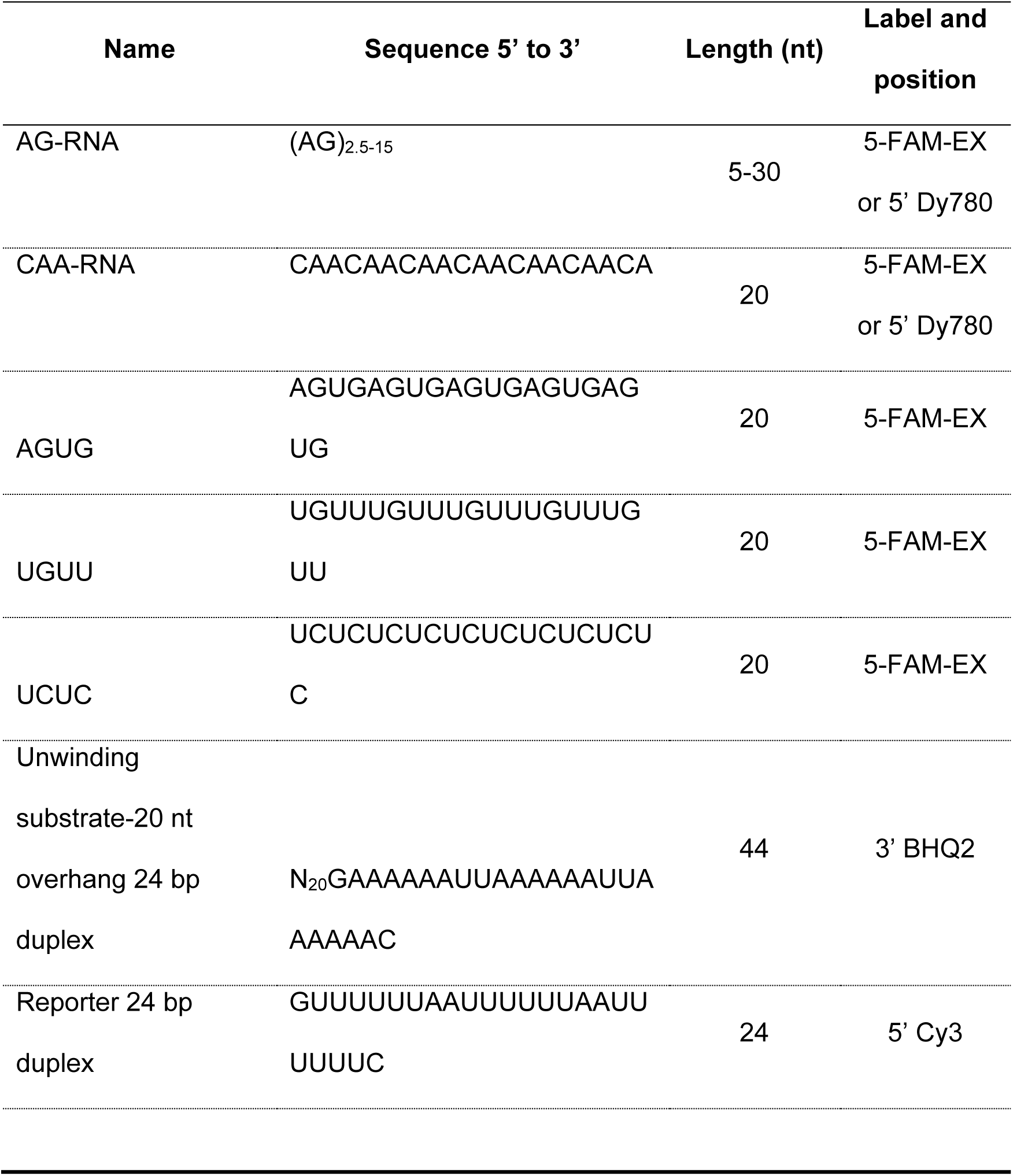
RNAs used in this study.

**Supplementary Table 2.**
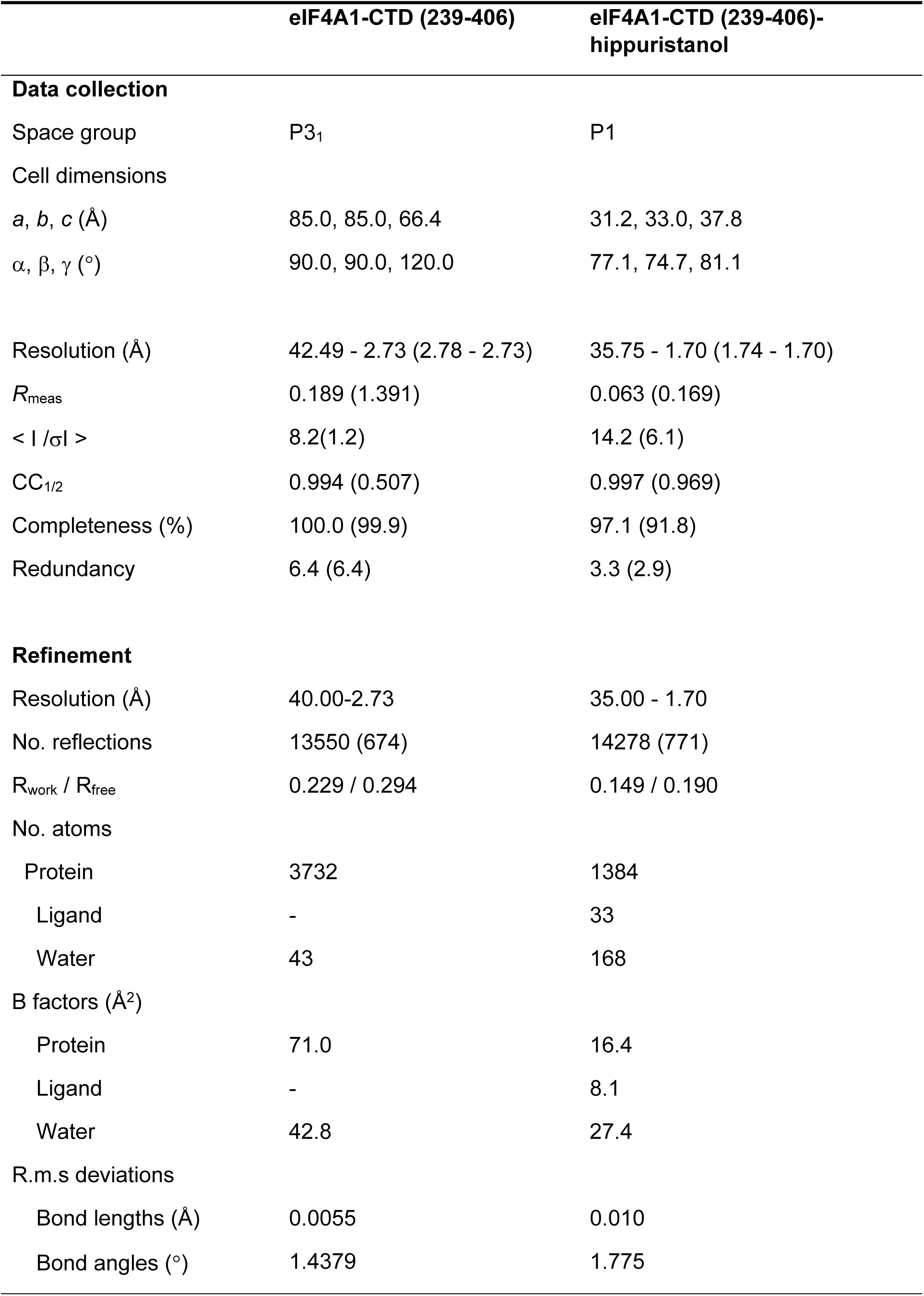
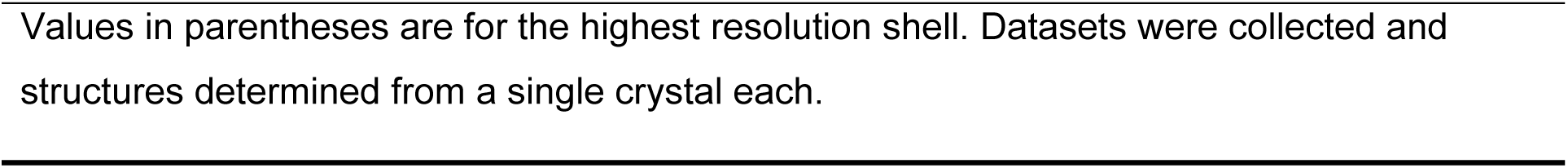
X-ray data collection and structure refinement.

|  | eIF4A1-CTD (239-406) | eIF4A1-CTD (239-406)-<br>hippuristanol |
| --- | --- | --- |
| **Data collection** |  |  |
| Space group | P3<sub>1</sub> | P1 |
| Cell dimensions |  |  |
| *a*, *b*, *c* (Å) | 85.0, 85.0, 66.4 | 31.2, 33.0, 37.8 |
| $\alpha$, $\beta$, $\gamma$ (°) | 90.0, 90.0, 120.0 | 77.1, 74.7, 81.1 |
| Resolution (Å) | 42.49 - 2.73 (2.78 - 2.73) | 35.75 - 1.70 (1.74 - 1.70) |
| *R*<sub>meas</sub> | 0.189 (1.391) | 0.063 (0.169) |
| $\langle I / \sigma I \rangle$ | 8.2(1.2) | 14.2 (6.1) |
| CC<sub>1/2</sub> | 0.994 (0.507) | 0.997 (0.969) |
| Completeness (%) | 100.0 (99.9) | 97.1 (91.8) |
| Redundancy | 6.4 (6.4) | 3.3 (2.9) |
| **Refinement** |  |  |
| Resolution (Å) | 40.00-2.73 | 35.00 - 1.70 |
| No. reflections | 13550 (674) | 14278 (771) |
| *R*<sub>work</sub> / *R*<sub>free</sub> | 0.229 / 0.294 | 0.149 / 0.190 |
| No. atoms |  |  |
| Protein | 3732 | 1384 |
| Ligand | - | 33 |
| Water | 43 | 168 |
| B factors (Å<sup>2</sup>) |  |  |
| Protein | 71.0 | 16.4 |
| Ligand | - | 8.1 |
| Water | 42.8 | 27.4 |
| R.m.s deviations |  |  |
| Bond lengths (Å) | 0.0055 | 0.010 |
| Bond angles (°) | 1.4379 | 1.775 |

Values in parentheses are for the highest resolution shell. Datasets were collected and structures determined from a single crystal each.

**Supplementary Table 3.**
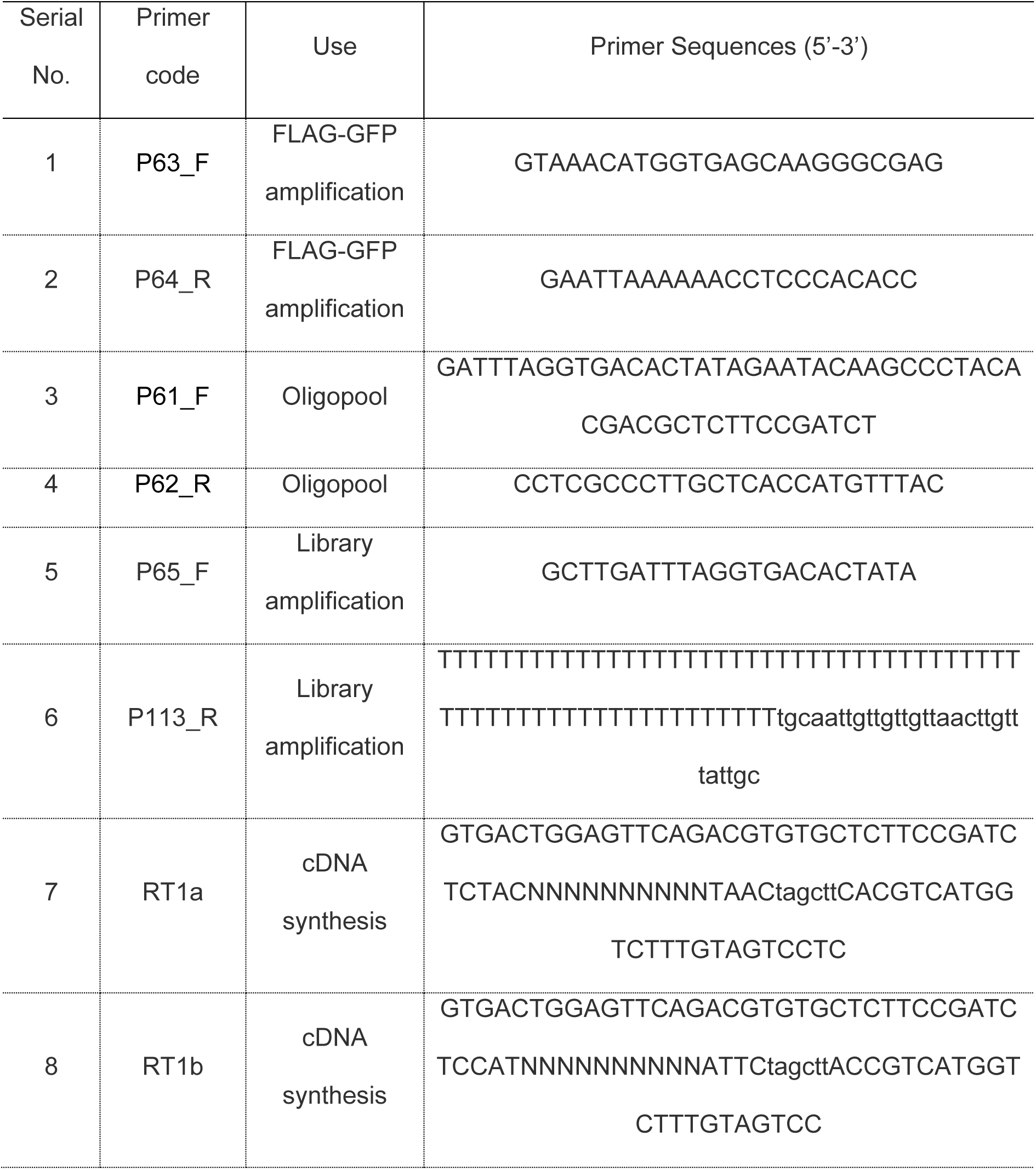
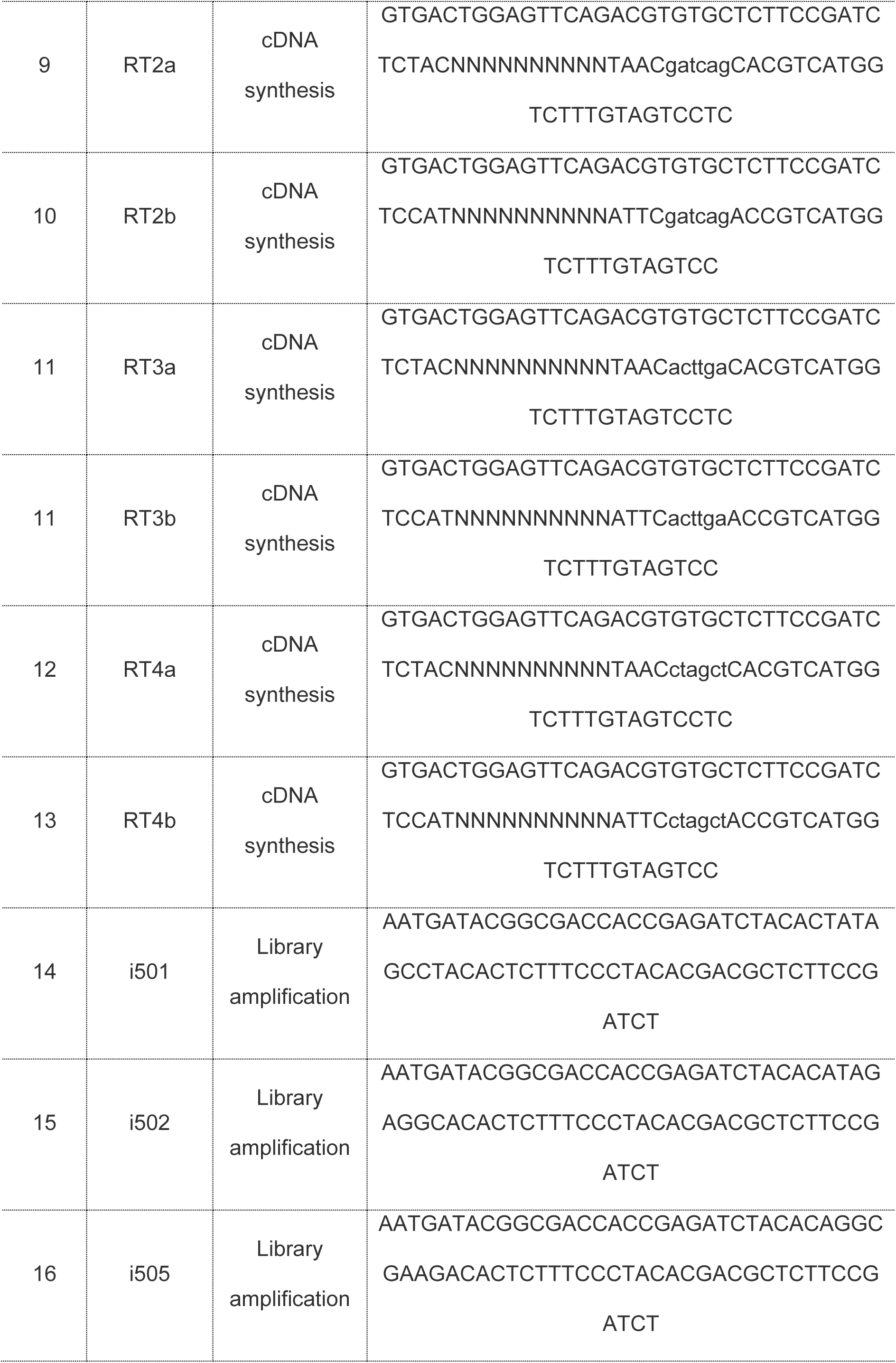
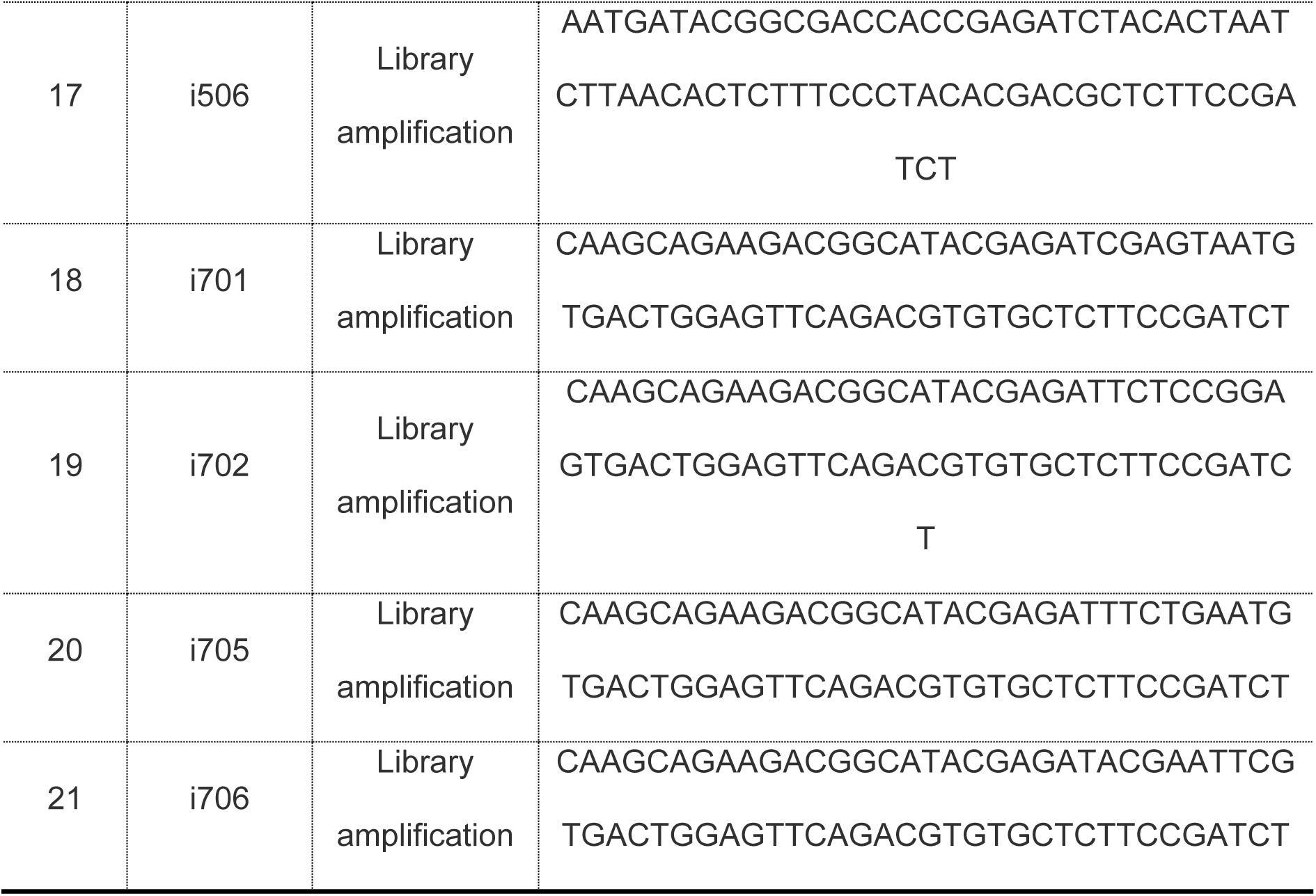
Oligonucleotides used for NaP-TRAP experiments.

